# Lowering the Thermal Noise Barrier in Functional Brain Mapping with Magnetic Resonance Imaging

**DOI:** 10.1101/2020.11.04.368357

**Authors:** L. Vizioli, S. Moeller, L. Dowdle, M. Akçakaya, F. De Martino, E. Yacoub, K. Ugurbil

## Abstract

Functional magnetic resonance imaging (fMRI) has become one of the most powerful tools for investigating the human brain. However, virtually all fMRI studies have relatively poor signal-to-noise ratio (SNR). Here we introduce a novel fMRI denoising technique, which suppresses noise that is indistinguishable from zero-mean, Gaussian-distributed noise. Thermal noise, falling in this category, is a major source of noise in fMRI, particularly, but not exclusively, at high spatial and/or temporal resolutions. Using 7-Tesla high-resolution data, we demonstrate improvements in temporal-SNR, the detection of stimulus-induced signal changes, and functional maps, while leaving stimulus-induced signal change amplitudes, image spatial precision, and functional point-spread-function unaltered. We also show that the method is equally applicable when using supra-millimeter resolution 3- and 7-Tesla fMRI data, different cortical regions, stimulation/task paradigms, and acquisition strategies. This denoising approach improves key metrics of functional activation detection while preserving spatial precision.

## Introduction

Since its introduction in 1992, functional Magnetic Resonance Imaging (fMRI)^1–3^ based on Blood Oxygenation Level Dependent (BOLD) contrast evolved to become an indispensable tool in the armamentarium of techniques employed for investigating human brain activity and functional connectivity. As such, it has been the central approach engaged in major initiatives targeting the human brain, such as the Human Connectome Project (HCP)^4^, UK Biobank project^5^, and the BRAIN Initiative^6^.

In all techniques employed in imaging biological tissues, the need for improving the spatiotemporal resolution is self-evident and, fMRI is no exception. To date, this challenge has been addressed primarily by increasing the magnetic field strength, leading to the development of ultrahigh magnetic field (UHF) of 7 Tesla (7T)^7^. UHF increases both the intrinsic signal-to-noise ratio (SNR) of the MR measurement as well as the magnitude and the spatial fidelity (relative to neuronal activity) of the BOLD based functional images^7–9^. These UHF advantages have enabled fMRI studies with submillimeter resolutions in the human brain, leading to functional mapping of cortical columns and layers, and other fine-scale organizations^7–9^. Such studies provide unique opportunities for investigating the organizing principles of the human cortex at the mesoscopic scale, thus bridging the gap between invasive electrophysiology and optical imaging studies and non-invasive human neuroimaging.

Despite these successes, however, the signal-to-noise and the functional contrast-to-noise ratios (SNR and fCNR, respectively) of fMRI measurements remain relatively low. This represents a major impediment to expanding the spatiotemporal scale of fMRI applications as well as the utility, interpretation, and ultimate impact of fMRI data.

What is considered “noise” in an fMRI time series is a complex question. Thermal noise associated with the MR detection^10, 11^, arising either from the electronics and/or the sample, is an important noise source in fMRI and would classify as a zero-mean Gaussian distributed noise. The use of parallel imaging to accelerate image acquisition, as is commonly done in contemporary MR imaging, introduces a spatially non-uniform amplification of this “thermal” noise by the g-factor^12^. The conditions under which this noise becomes dominant in an fMRI time series depends on the static magnetic field strength, the voxel volume, and image repetition time (TR) used in the experiment, becoming more prominent at higher resolutions (i.e. smaller voxel volumes), short TRs, and/or lower magnetic fields^13, 14^. It is the *dominant* contribution at ∼0.5 µL voxel volumes (e.g. ∼0.8 mm isotropic dimensions) typically employed in high resolution 7T fMRI studies; it remains dominant at 7T up to ∼10 µL voxel volumes, gradually plateauing beyond that^13, 14^. However, even with 3 mm isotropic resolution (i.e. 27 µL voxel volume) and relatively long TR acquisitions, thermal noise was estimated to be a significant contributor to fMRI time series at 7T^15^. At lower magnetic fields like 3T, where this type of noise becomes more conspicuous, and where typical fMRI resolutions employed are ≲3 mm, it would be a substantial contributor in virtually all fMRI studies^13, 14^.

In this paper, we tackle these SNR and fCNR limitations using a denoising technique – namely, Noise Reduction with Distribution Corrected (NORDIC) PCA. NORDIC operates on repetitively acquired MRI data and only removes components which cannot be distinguished from zero-mean Gaussian distributed noise; as such, the method targets the suppression of thermal noise and not the structured, non-white noise caused by respiration, cardiac pulsation, and spontaneous neuronal activity (e.g.^16–19^ and references therein).

High resolution 7 Tesla data, as well as data obtained with more conventional, supra-millimeter resolution at 3T and 7T using several different task/stimulus and acquisition strategies, demonstrate that major gains are achievable under a wide variety of experimental conditions with NORDIC in gradient-echo (GE) BOLD fMRI without introducing image blurring. Based on these findings, the approach is expected to markedly widen the scope and applications of fMRI in general, and high spatial and/or temporal resolution fMRI in particular.

## Results

The fMRI data, acquired with GE Simultaneous Multi Slice (SMS)/Multiband (MB) Echo Planar Imaging (EPI)^20, 21^, were reconstructed either by the algorithms provided with the MR scanner (referred throughout this work as “Standard”), or by the NORDIC PCA method (see Methods for a detailed description) using the raw k-space files produced by the scanner (referred to as “NORDIC”).

The bulk of the analyses were performed on data acquired on 4 subjects with a variant of a widely used, 0.8 mm isotropic resolution 7T protocol (see Methods) based on a block design visual stimulation paradigm (Figure 1A); these analyses are presented in this section. However, to ensure the generalizability our results and the versatility of the NORDIC approach, we present as Supplementary Material evaluations of NORDIC on fMRI across acquisition parameters, field strengths (i.e. 3T and 7T), cortical regions, and stimulation paradigms, bringing the total number of datasets to N = 10. All data sets showed converging results.

**Figure 1.**
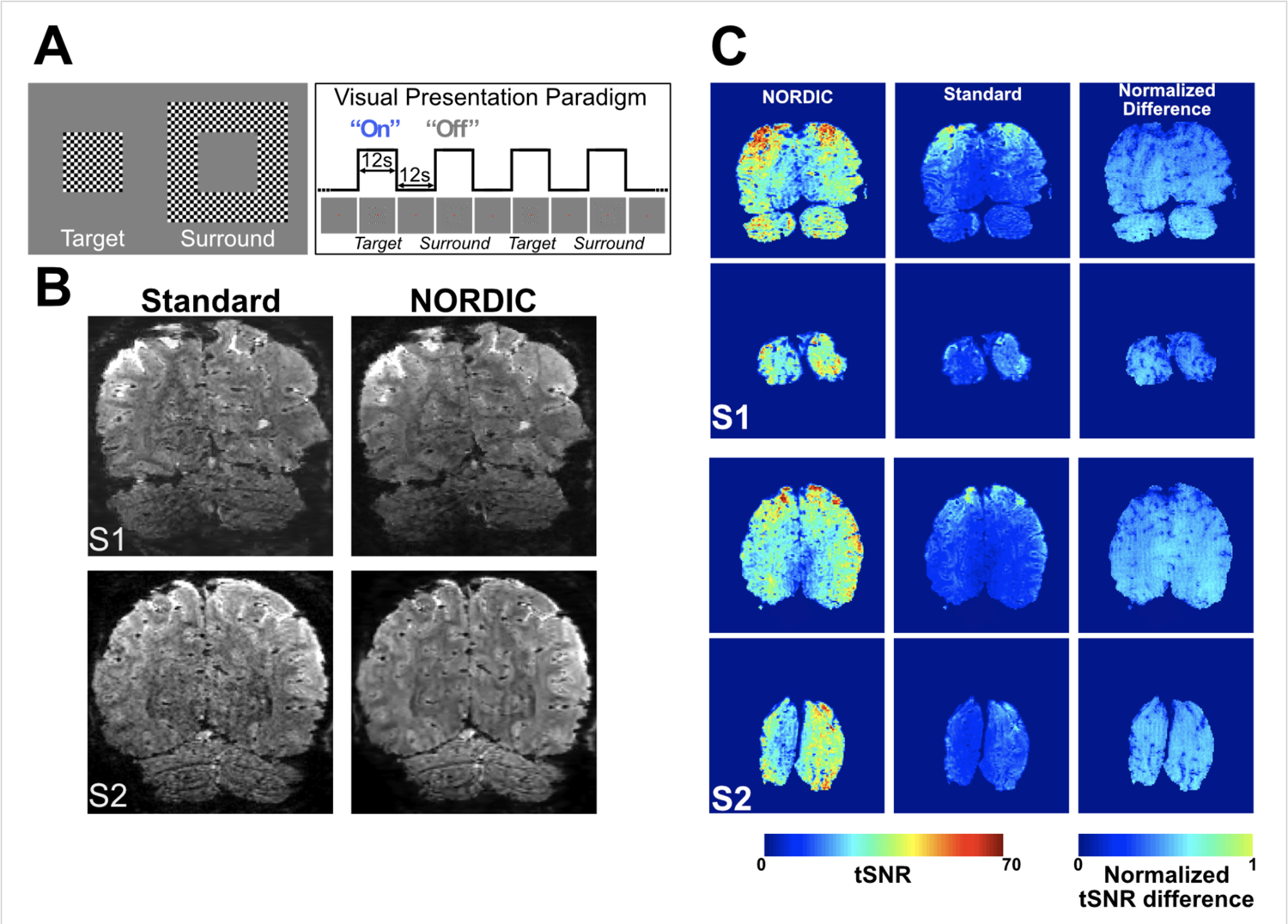
Stimuli and paradigm, epi images and tSNR. Panel A depicts the visual stimuli (left) used and a schematic of the visual presentation paradigm (right). Panel B shows an example slice from a single volume extracted from an fMRI time series for Standard (left column) and NORDIC (right column) reconstructions before any preprocessing, for two subjects S1 and S2. Panel C shows average (across all 8 runs) brain temporal signal-to-noise ratio (tSNR) maps of 2 exemplar slices in 2 representative subjects (S1 and S2) for NORDIC (left) and Standard (center) reconstructions and the normalized difference between the 2. The last was computed by performing (tSNR_NORDIC_ - tSNR_STANDARD_)/tSNR_NORDIC_). The slices chosen represent one of the anterior most slices in the covered volume, and an occipital slice that includes a portion of the target ROI in V1.

We used a block design, visual stimulation paradigm comparable to that implemented in Shmuel et al.^22^ with minor modifications: It consisted of retinotopically organized target and surround stimuli presented in alternating stimulus-on and -off epochs (Figure 1A). Each “run” consisted of six stimulus-on epochs, three each for target and surround stimuli. We acquired 8 experimental runs in 6 subjects (4 at 7T and 2 at 3T, the latter presented as Supplementary Material); 2 of these runs were used to identify the retinotopic representation of the target in V1, computed by contrasting the target *versus* the surround condition (p<0.01 uncorrected). This functionally defined region of interest (ROI), referred to as “target ROI” from here on, was subsequently used for all ROI confined analyses. The functional runs used to estimate the ROI were excluded from subsequent analyses (see Methods).

### NORDIC vs. Standard MR Images

Figure 1B illustrates an example slice for Standard and NORDIC reconstructed GE-EPI images for two subjects before any preprocessing for fMRI analysis was applied. An improvement is visually perceptible for NORDIC images, especially in the central regions where the g-factor noise amplification would be particularly elevated (see also Figure 6A). Subtraction of Standard from NORDIC processed image of a single slice from a single timepoint in the fMRI times series displayed only noise without any features of the image or edge effects; when such a difference was calculated for all time points in the fMRI time series and averaged, the result was equivalent to the g-factor map (Supplementary Figure 1). These observations are consistent with NORDIC suppressing only random noise without impacting the image.

Figure 1C shows temporal SNR (tSNR) maps averaged across all eight runs for two exemplar subjects and slices. The average tSNR across *all* the voxels in the brain was more than 2-fold larger for NORDIC (S1_tSNR_: 27.34±2.26 (std); S2_tSNR_: 33.01±2.26 (std); S3_tSNR_: 43.31±1.52 (std); S4_tSNR_: 26.61±2.32 (std)) compared to Standard (S1_tSNR_: 13.28±0.19 (std); S2_tSNR_: 13.97±0.05 (std); S3_tSNR_: 16.1±0.26 (std); S4_tSNR_: 14.12±0.23 (std)) images. Paired sample t-tests carried out across all 8 runs, independently per subject, indicated that for all subjects, the average tSNR for NORDIC was significantly larger (p<0.01e^-5^) than that for Standard images. Improvements in tSNR with NORDIC in individual runs are shown in Supplementary Figures 2 and 3.

### Functional Images

Impact of NORDIC on functional maps was evaluated by comparing a single run processed with NORDIC against the concatenation of multiple runs of the Standard reconstruction (see Methods). Figure 2 illustrates functional maps on the inflated surface of one hemisphere, contrasting the target *versus* the surround condition thresholded at t≥|5.7| for four subjects. For two subjects, representative single run functional maps are also shown for two different t-thresholds and on anatomical image of a slice in Supplementary Figure 4. At the same t-threshold, the extent of activation achievable with a single NORDIC run was comparable or better than that obtained by concatenating 3 to 5 Standard runs.

**Figure 2.**
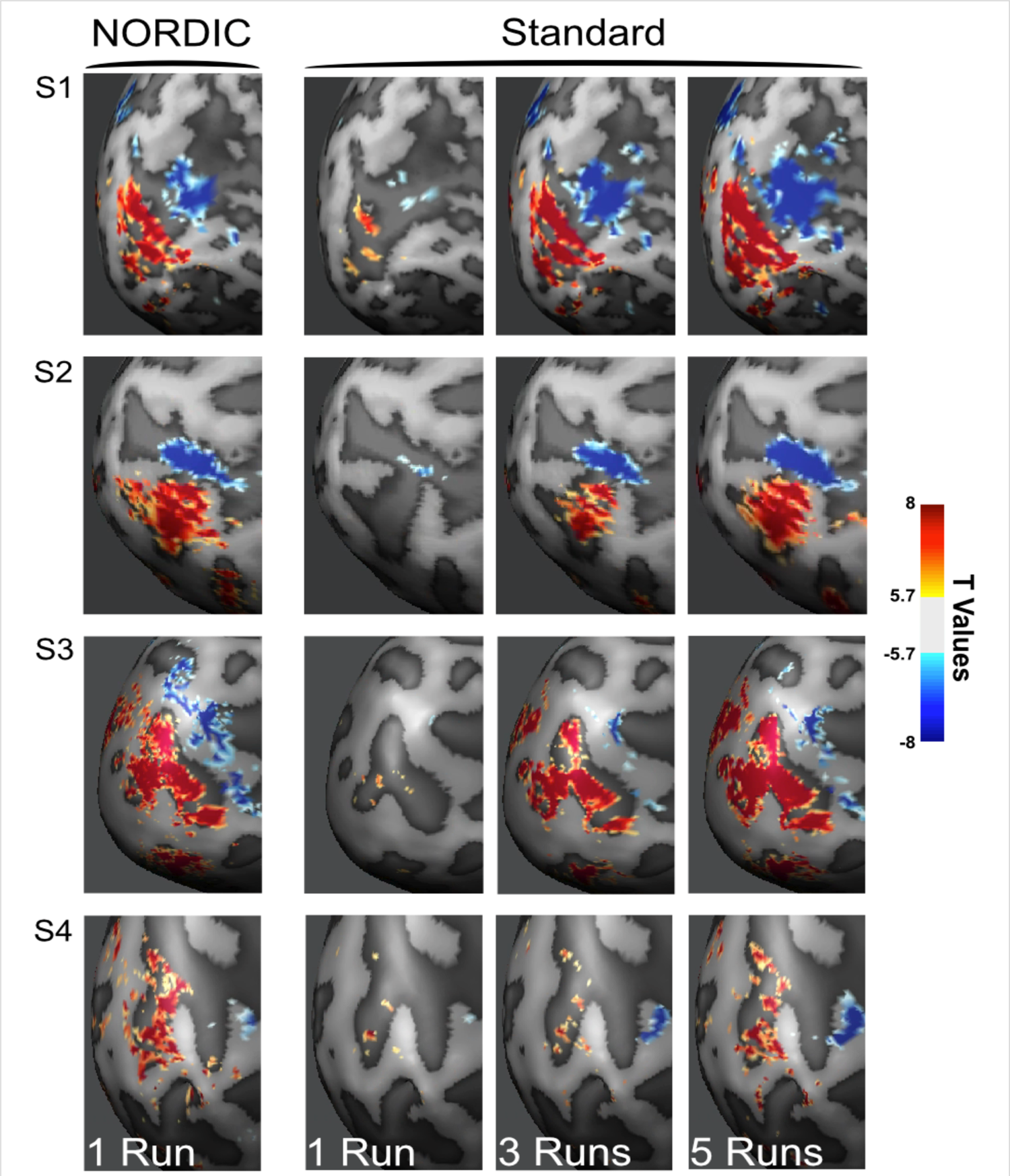
NORDIC vs. Standard t-Maps. Left most panel shows functional images as t-maps (target > surround) thresholded at t ≥ |5.7| for a single NORDIC processed run, and for 1, 3 and 5 Standard processed runs combined, for subject 1 (S1), subject 2 (S2), subject 3 (S3) and subject 4 (S4).

Similar results are presented in Supplementary Materials for supra-millimeter 3T and 7T fMRI data obtained with visual stimulation and face recognition paradigms (Supplementary Figures 8-11), and for 0.8 mm 7T data obtained with auditory stimulation (Supplementary Figure 12); two of these datasets (Supplementary Figures 10, 11) were acquired with an event related paradigm.

Consistent with the data displayed in Figure 2, the t-values examined further in two subjects (S1 and S2) were significantly larger for NORDIC (p<0.05) than its Standard counterpart (Figure 3A and 3E) within the target ROI, as determined with linear mixed models carried out independently per subject. When the t-value distribution for the target>0 contrast was analyzed for three ROIs (Supplementary Figure 6), it was found to be shifted to higher values in each individual run for the target ROI; for the two other ROIs in regions where stimulus evoked responses should not exist, it was essentially unaltered, demonstrating that NORDIC does not perturb t-values where it should not.

**Figure 3.**
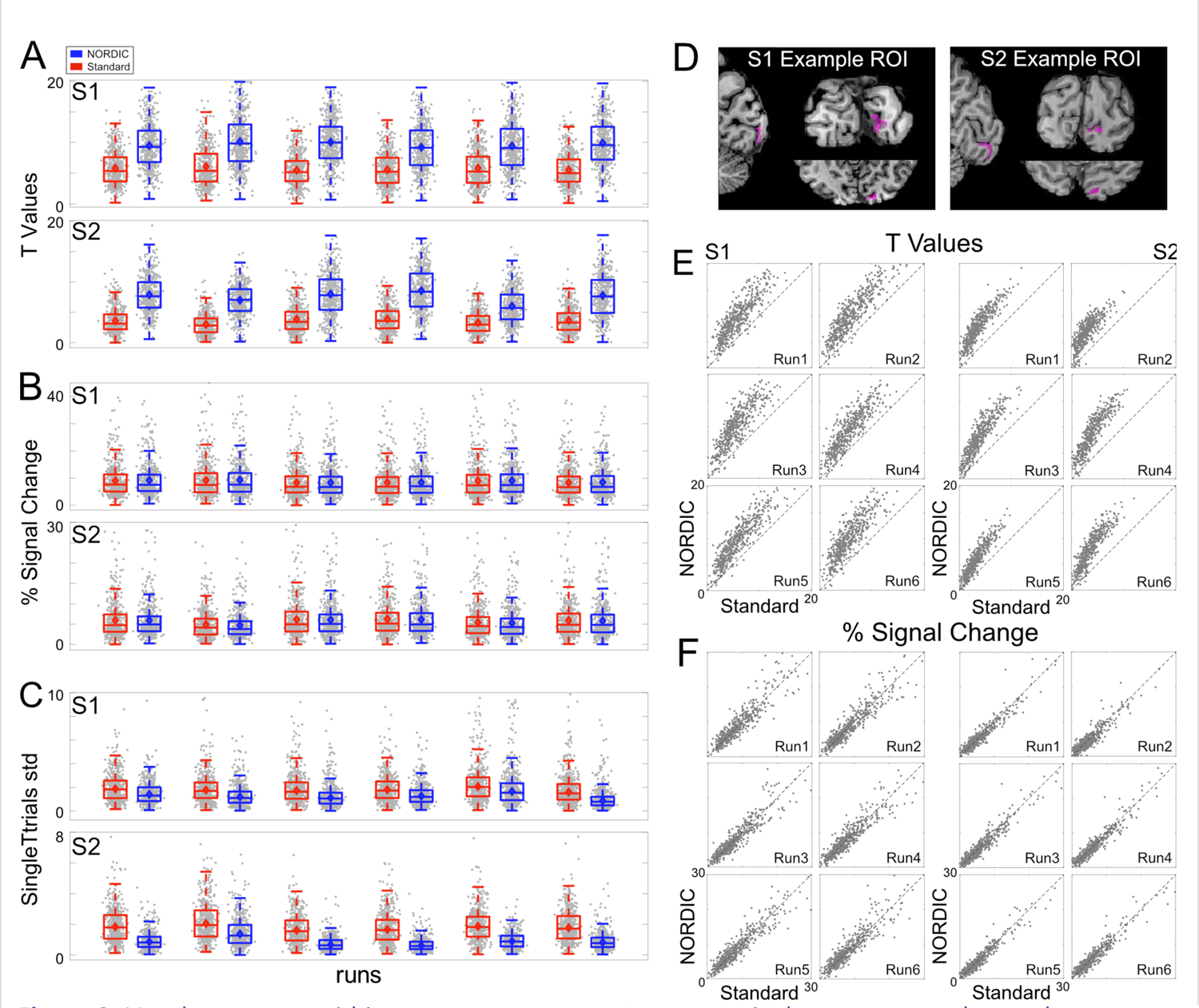
Voxel responses within target ROI. Panel A shows the single-run (arranged over the x axis) t-values (activity elicited by the target > 0) induced by the target stimulus for standard (red) and NORDIC (blue) data. Panel B is the same as panel A, but for beta weights (transformed into percent signal change). Panel C shows the single-run standard deviation computed across single trial PSC beta estimates elicited by the target condition. For these 3 panels, grey dots represent responses to single voxels with the target ROI (497 for S1 and 461 for S2). The box-and-whisker plots, computed across all ROI voxels, represent the interquartile range (IQR – with box limits being the upper and lower quartile), with the whiskers extending 1.5 time the IQR or to the largest value. The horizontal lines within the boxplot represent the median, while the diamond the mean across voxels. Panel D shows the target ROI, representing the left retinotopic representation of the target in V1 for 2 exemplar subjects in all 3 planes. Panel E shows the single runs, single voxel scatterplots for t-values (activity elicited by the target > 0), for Standard (x axis) and NORDIC (y axis). Panel F: same as panel E for the beta percent signal change responses to the target condition.

Percent signal change (PSC) within the target ROI as the mean of all voxels and at the single-voxel level are presented in Figures 3B and 3F, respectively. The stimulus-induced PSC was highly comparable across reconstruction types; linear mixed models carried out independently per subject (with the individual runs as random effect) showed no significant (p>0.05, Bonferroni corrected) differences in PSC amplitudes across reconstructions for all runs.

Figure 3C depicts the standard deviation (20% trimmed mean across voxels within target ROI^23^) computed amongst PSC betas elicited by a single presentation of the target stimulus within a run. As shown by both paired sample t-test (p<0.05) and 95% bootstrap confidence interval (carried out by sampling with replacement of the individual runs), this metric was found to be significantly larger for Standard than NORDIC, indicating greater stability of NORDIC PSC single trial estimates among the different stimulus epochs within a run.

The equivalency of PSC amplitudes for NORDIC and Standard reconstructions are further illustrated using images in Figure 4 and Supplementary Figure 5. In addition, a hold-out data analysis was carried out with PSC estimates (Figure 4); for this, we estimated GLM model parameters in one run and assessed the precision with which these parameters predicted the PSC in all other runs at a single voxel level. The precision of PSC estimates, computed as cross-validated R2 for single run GLMs was higher (Figure 4A, third row) for the NORDIC compared to the Standard reconstruction. Paired sample 2-sided t-test carried out across cross-validations folds showed that within the target ROI average R2 (see Methods) was significantly (p<0.01 Bonferroni corrected) higher for NORDIC (S1: NORDIC mean R2=36.43 (ste=9.81); Standard mean R2=22.2 (ste=5); S2: NORDIC mean R2=25.36 (ste=5.74); Standard: mean R2=10.62 (ste=3.2) and Figure 4B, bar graphs), indicating again higher precision of PSC estimates and their stronger predictive value for NORDIC.

**Figure 4:**
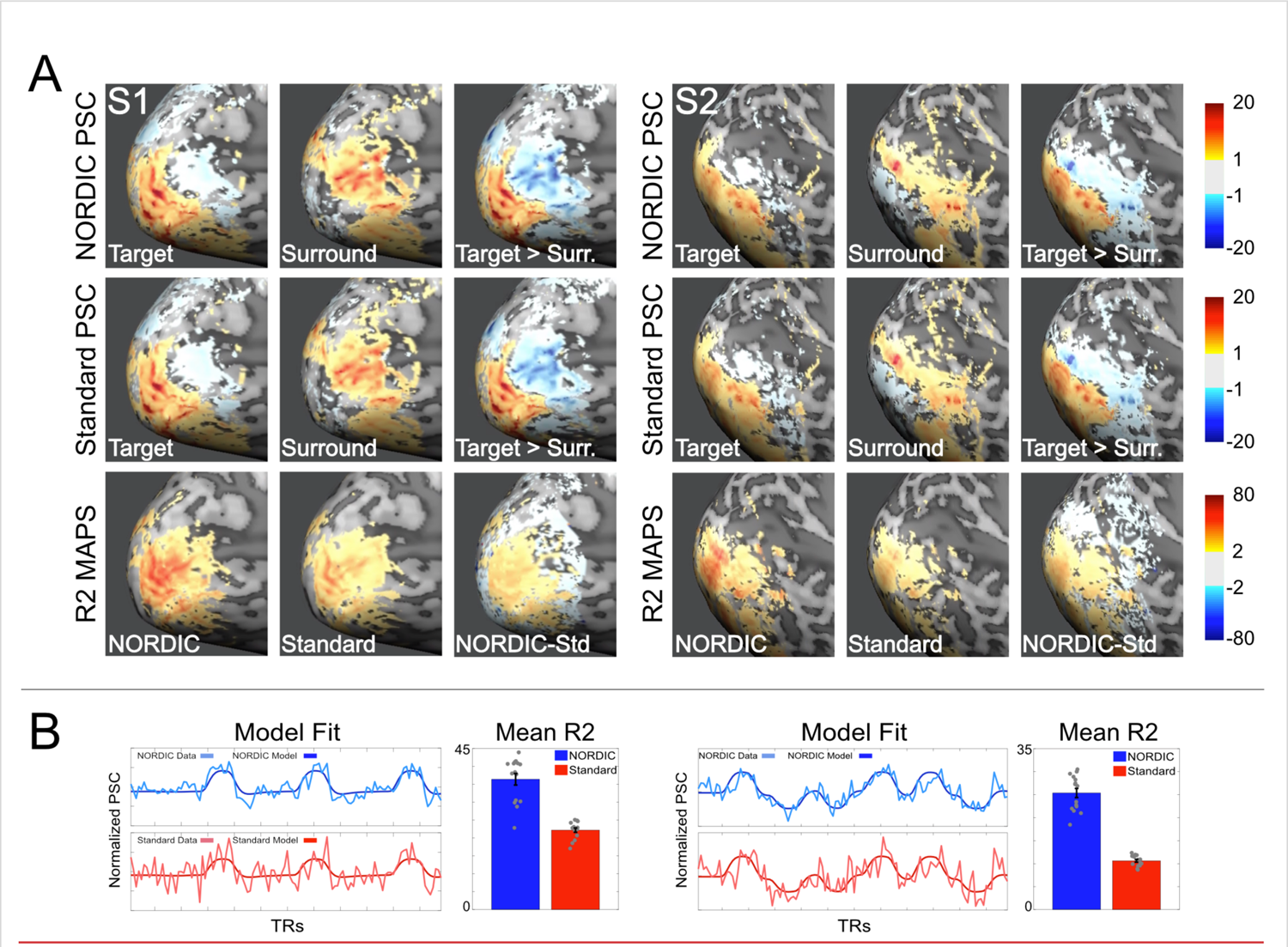
PSC maps and cross-validated prediction accuracy. Panel A: Average percent signal change (PSC) maps and cross-validated R2 for both subjects S1 and S2. The top 2 rows in Panel A show the average (across runs) PSC maps elicited by the target (left), surround (middle) and their contrast target>surround (right), for NORDIC and Standard reconstructions, respectively. As evident by these images, PSC amplitude and the extent of stimulus induced signal change is comparable across reconstructions (see also Supplemental Figure 5). The 3^rd^ row in panel A shows the average (across folds) cross-validated R2 maps for NORDIC (left), Standard (middle) and their difference (right). *Only in the relevant portion of the cortex for the stimulus used* (i.e. areas where stimulus induced BOLD activity is expected, as indicated by the PSC maps) the R2 maps show higher precision of PSC estimates for NORDIC images. Panel B: For each subject, left column show a single run of NORDIC and Standard BOLD time-courses for a target-selective voxel (lighter lines) and its prediction estimated on a separate run (darker lines). These plots highlight a closer correspondence between model prediction and empirical time-courses for NORDIC data, as summarized by the significantly larger (2-sided paired sample t-test; S1: t(14)=12.8, ci(11.83;16.6), Cohen’s d: 3.31 p<.01 Bonferroni corrected; S2: t(14)=20.1, ci(13.33;16.25), Cohen’s d: 5.41 p<.01 Bonferroni corrected) target ROI average cross-validate R2 (bar plots represent the mean and error bars indicate standard errors of the mean across 15 cross-validation folds shown as grey dots).

Figure 5 shows the functional point spread (PSF) measurements on the cortical surface calculated following previous work^22^ using NORDIC and Standard images (see Methods) from two subjects: briefly, the approach defines the boundary between the target and the surround stimuli as those voxels showing a *differential* functional response close to 0 (Figure 5A, left column for each subject). Along traces drawn orthogonal to this boundary, the functional response amplitudes are then measured in the *single condition* maps and subsequently quantified by fitting a model consisting of a step-function (representing infinitely sharp PSF) convolved with a Gaussian^22^ (Figure 5B). The full width half max (FWHM) of the Gaussian represents the functional PSF^22^. With NORDIC, the average PSFs (across traces) were 1.04 mm (std: 0.19) and 1.22 mm (std: 0.51), for subjects 1 and 2, respectively; the average PSFs for the Standard were 1.14 mm (std: 0.16) and 1.15 mm (std: 0.11). Paired sample t-tests carried across the 8 runs showed no significant differences (p>0.05) in functional PSF amongst reconstruction types.

**Figure 5.**
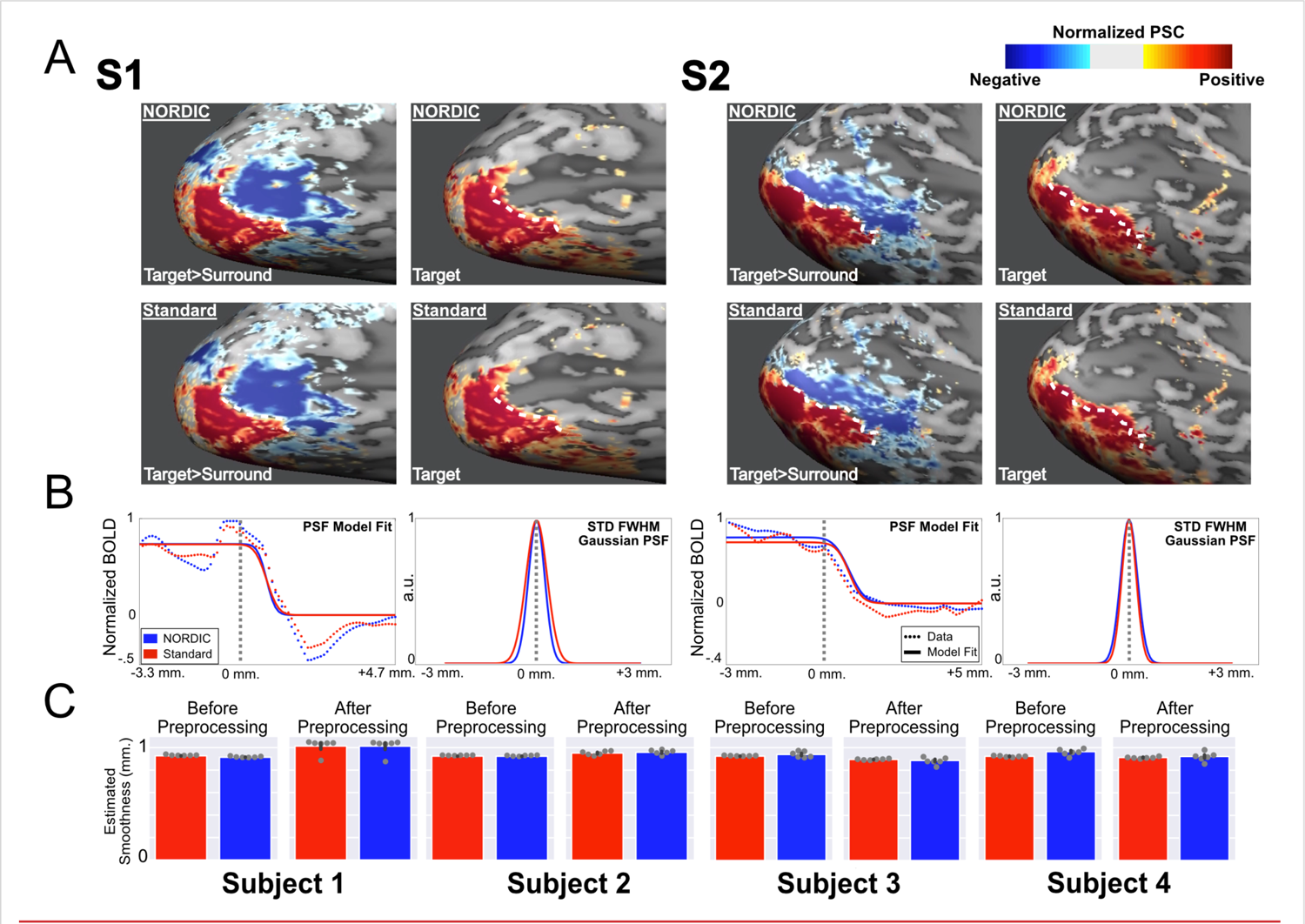
Functional point spread function (PSF) and global image smoothness. Panel A, top row shows NORDIC normalized beta percent signal change (PSC) maps for *differential* mapping target (in red) > surround (in blue) (left), and the target only (right) *single-condition* image for subjects 1 and 2. The white dotted line is determined in the *differential* image as the “boundary” between the two stimulations. The same white dotted line is also superimposed on the target only PSC map where PSC values are greater than zero but decreasing in magnitude progressively away from this “boundary” posteriorly. The functional PSF is calculated from this spread in PSC beyond the “boundary”. Panel A, lower row is identical to the upper row, but obtained from Standard reconstruction data. Panel B, left panel for each subject: the PSC magnitude changes (normalized to the highest value) along traces perpendicular to the “boundary” are displayed as the average (across traces and runs). The model fits (solid line) and data (dotted line) are shown for both the NORDIC (blue) and Standard (red) reconstructions. The vertical grey dotted line represents the “boundary” as derived from the differential maps. Panel B, right panel for each subject portrays the full width half max (FWHM) standard deviation of the gaussian kernel that was convolved with a step function to model functional PSF (see Methods). Panel C: Mean global smoothness of images used for the fMRI time series for Standard (red) and NORDIC (blue) in four subjects, before (left panel) and after preprocessing related interpolations. Error bars represent standard error of the mean across 6 independent runs (shown as grey dots).

**Figure 6.**
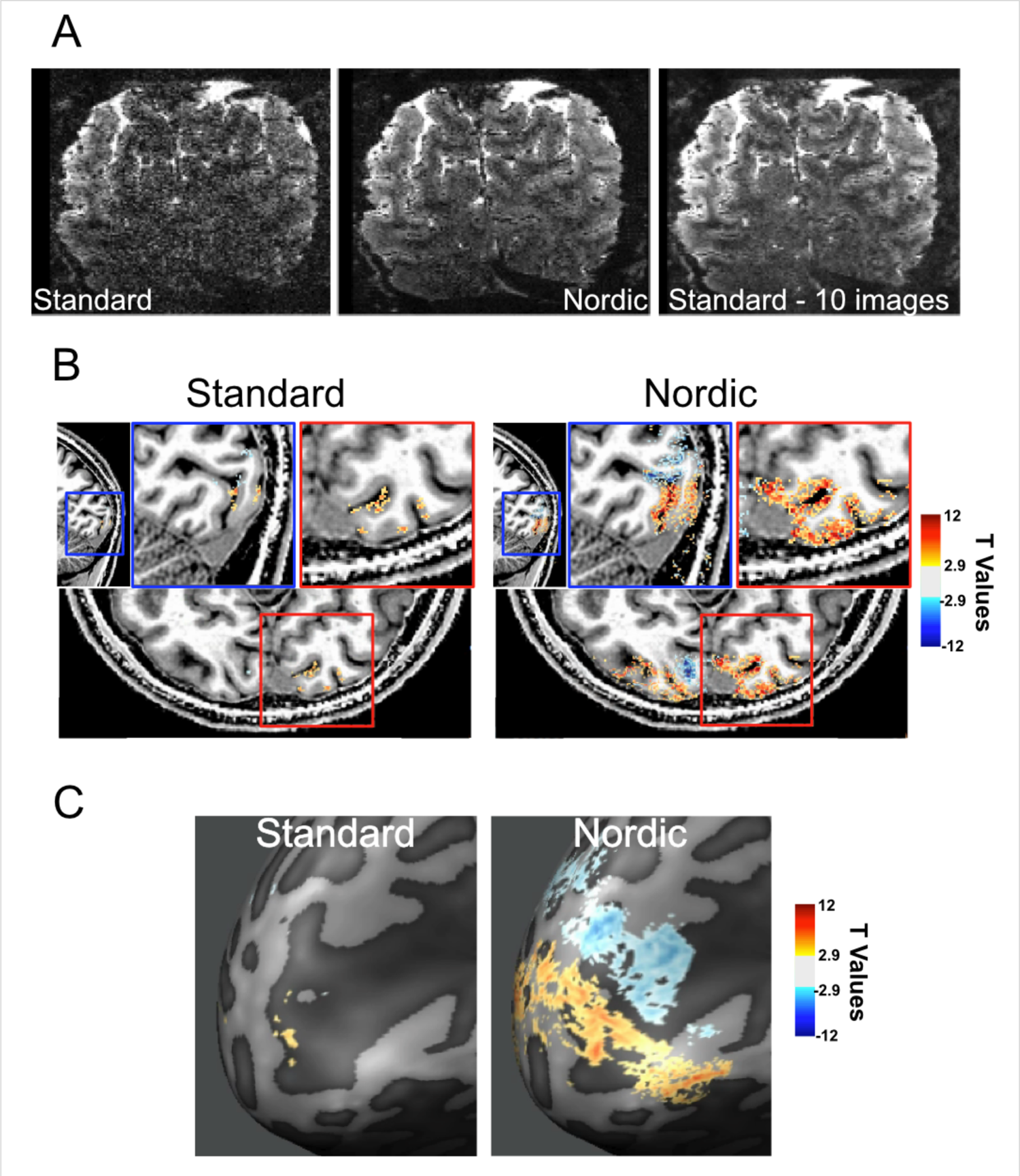
3D GE EPI images and fMRI data obtained with 0.5 mm isotropic voxels. Panel A shows a single slice from a single time point in the consecutively acquired volumes forming the fMRI time series for Standard (left) and NORDIC (middle) images. The right panel shows the average of 10 images of the same slice for the Standard reconstruction. Panel B shows t-thresholded (|t| ≥ 2.9) functional maps (for the contrast target>surround on a T1-weighted anatommical image for standard (left) and NORDIC (right) reconstructions for a saggital and axial slice (with related zoom-ins on the sagittal (blue) and axial (red) planes). Panel C shows the same t-maps as in panel C on the inflated cortical space and at different t-thresolds. No spatial smoothing or masking was applied to the data.

In addition, we estimated the global smoothness of individual GE-SMS/MB-EPI images in the fMRI time series using AFNI (3dFWHMx function)^24^, with automatic intensity-based masking derived from the median image of each run. The spatial autocorrelation was estimated using a Gaussian+monoexponential decay mixed model to account for possible long-tail autocorrelations. The FWHM from this mixed model estimate, averaged over 4 subjects, before and after data preprocessing (see Methods) for the Standard reconstruction was 0.92±0.002mm, and 0.94±0.05mm, respectively; for the NORDIC reconstruction, these values were 0.93±0.02 and 0.94±0.05 mm (Figure 5C). A linear mixed model carried out across subjects and runs indicated non meaningful differences in smoothness estimate (p>0.05) between Standard and NORDIC.

Figure 6 shows 0.5 mm isotropic resolution fMRI data (0.125 µL voxel volume) obtained using the target/surround visual stimulation paradigm (see Methods and also Supplementary Figure 14). Figure 6A and Supplementary Figure 14A display a single coronal slice in the visual cortex from one of the repetitively acquired volumes in the fMRI time series. Processed with the Standard reconstruction, the image of this slice is noisy and practically unusable for functional mapping. However, the single image after NORDIC reconstruction and average of 10 images from the Standard reconstruction look virtually identical; these very high-resolution data also demonstrate clearly that NORDIC does not induce smoothing (see expanded panels in Supplementary Figure 14 and also Panel D in Supplementary Figure 16).

Functional maps from the 0.5mm data computed for the target > surround contrast using the 8 concatenated runs (i.e. ∼44 minutes of data for the two stimulus conditions and interleving baseline periods) are shown superimposed on T_1_-weighted anatomical images (Figure 6B) and the flattened cortex (Figure 6C). These functional data do not have any spatial smoothing or masking applied to them. Little activation is detected with Standard reconstruction. Localized and highly precise BOLD activation, allowing differentitation of adjacent sulcus banks (Figure 6B) are observed for NORDIC images. Consistent with these observation, stimulus-evoked signal changes in the fMRI time course at a single voxel level were virtually undetectable in Standard reconstruction but obviously visible with the use of NORDIC (Supplementary Figure 14B).

The NORDIC method should be equally applicable for resting state (rsfMRI) that is used extensively to evaluate functional connectivity. We present a preliminary analysis on one subject at 3T confirming this expectation (Supplementary Figure 13).

As previously mentioned, the “Standard” reconstruction employed in these comparisons is the one provided by the vendor of the scanner. For NORDIC, prior to denoising, the same experimentally acquired k-space data was exported and had to be processed “offline” for EPI and GRAPPA reconstructions using our own implementation. We had opted to use the vendor provided reconstruction for the comparisons because we wanted to demonstrate improvements attainable with NORDIC relative to what is available for the general fMRI community, which relies on reconstruction provided by the vendor of their scanners, in this case Siemens scanners. This choice, however, raises the question of whether the gains demonstrated are not only due to NORDIC but are also partially related to differences in the reconstruction pipeline. To address this question, we have reproduced Figure 2 for one of the four subjects (subject S2) using our offline reconstruction pipeline but *without* the NORDIC denoising step. Supplementary Figure 15 displays the results both for “Scanner Standard”, which, for ease of comparison, duplicates the functional images shown in Figure 2, and those obtained using our offline reconstruction (“Offline Standard”). The results demonstrate that in the three cases shown (1 run, and 3 and 5 runs concatenated), the Scanner Standard and the Offline Standard produce virtually identical results, and in both cases, functional maps of 5 concatenated runs look essentially identical to a single NORDIC run.

Despite being fundamentally different from NORDIC, global SVD or PCA based methods (e.g. ^25, 26^) can also identify random noise components in a time series and thus, can in principle be used to selectively suppress its contribution. Therefore, a comparison of NORDIC against such an approach would be informative. On the other hand, performing a thorough comparison under all possible conditions is beyond the scope of this paper, the primary aim of which is to introduce NORDIC and showcase its versatility (i.e. working well both in high and low SNR, cyclic and event related paradigms, 3 and 7T, and combinations thereof). Nevertheless, we present here the results of comparing NORDIC to two other PCA based approaches using the 0.5 mm isotropic fMRI data where suppressing random thermal noise without incurring meaningful spatial smoothing is most challenging and at the same time, of utmost importance. Supplementary Figure 16 illustrates these comparisons. This figure demonstrates that relative to a global PCA approach with a “white noise” criterion to identify random noise^26^ (labeled PCAwn), which essentially follows an earlier SVD approach^25^ with modifications, performance of NORDIC is far superior in terms of the individual images, t-statistics, spatial smoothing, and the resultant functional maps. This figure also contains a comparison to a widely available implementation (named DWIdenoise) for complex and magnitude data of a relatively recent denoising technique called Marchenko-Pastur Principle Component Analysis (MPPCA)^27^. Again, NORDIC outperforms this approach with respect to t-statistics and consequently t-thresholded functional maps, even though this method causes significantly larger smoothing than NORDIC.

The metrics presented in Supplementary Figure 16 are useful in evaluating the performance of different denoising algorithms when *taken together*. However, caution should be exercised in interpreting any one metric alone. For example, the smoothness estimate for PCAwn, taken alone, suggests that this method performs relatively well. However, if we examine the EPI images, t-value distribution, and the related functional map (Supplementary Figure 16), it becomes evident that this apparent preservation of spatial precision is an outcome of the failure to remove thermal noise. As explained earlier, smoothness metrics derived here utilize spatial autocorrelation, which is nonexistent for Gaussian zero-mean thermal noise. Images dominated by thermal noise would therefore show low smoothness estimates. Conversely, highly smoothed images would lead to low GLM residuals and therefore high t-values, albeit at the expense of degraded spatial precision.

## Discussion

fMRI is inherently a low contrast-to-noise measurement where the biologically driven responses are relatively small compared to fluctuations (i.e. “noise”) in the amplitude of the signal in the fMRI time series. Certainly, thermal noise of the MR detection^10, 11^ contributes to this “noise”. Physiological processes of respiration and cardiac pulsation^15, 28–30^, and for task/stimulus fMRI, the spatially correlated spontaneous fluctuations ascribed to functional networks in rsfMRI^31^ represent other sources of tSNR degradation, which, unlike thermal noise, are non-white in nature^16–19^. These non-Gaussian sources of signal fluctuations are proportional to signal magnitude^14, 32–35^; as such, they become dominant only when a voxel’s signal (which is proportional to voxel volume) is large compared to instrumental thermal noise, as encountered, for example, with low spatial resolutions, high flip angles used in conjunction with long TRs, and at high magnetic fields^13, 14, 36, 37^.

Reliably detecting the relatively weak biologically driven responses in the presence of the afore-mentioned noise contributions requires significant efforts to clean up the fMRI time series. This problem was addressed as early as approximately two decades ago using component analysis based on SVD^25^, and subsequently PCA and ICA^26^ to decompose the fMRI time series into components containing the task/stimulus response, structured noise, and thermal (random) noise. Although these early holistic approaches have not been widely adapted, numerous methods using PCA and ICA components analysis in various ways have subsequently been introduced and employed almost exclusively on the suppression of the non-white confounds (e.g. ^16–19, 38, 39^ and references therein).

In this paper, we introduce a method named NORDIC aimed at improving the detectability of the inherently small fMRI signals by selectively targeting the suppression of thermal noise. NORDIC is fundamentally different in its approach to the above referenced PCA and ICA methods. Although, these previous methods for the most part have concentrated on identifying structured noise^17^, some of them also provide a strategy to selectively suppress thermal noise; they do so using a global PCA analysis and an empirical threshold for the differentiation of noise and signal components, in some cases working best in the presence of a clear periodic temporal signature in the signal^26^, which naturally limits their general utility. In contrast, NORDIC uses a local (patch) approach, experimental characterization of thermal noise independent of the functional imaging data, and a well-defined objective principle to identify the threshold for its suppression. Especially for low SNR, high resolution fMRI data, a global component analysis may be suboptimal (see comparison in Supplementary Figure 16); as such, in such data, where the need for thermal noise suppression is immense, spatial smoothing has been the method of choice to improve SNR and fCNR even at the risk of degrading spatial specificity. In contrast, our results demonstrate that NORDIC is particularly (but not exclusively) useful for such low SNR high resolution fMRI data (Figure 6, Supplementary Figures 14, 16).

NORDIC and its application in diffusion weighted imaging (dMRI) was previously described^40^ and was shown to yield superior results to the recently introduced MPPCA^27^ method, which also selectively targets thermal noise removal. It is difficult to precisely identify the components that are removed in MPPCA, although its application leads to better results in dMRI^27, 40^ and increased reproducibility in rsfMRI^41, 42^. In contrast, NORDIC yields a parameter-free threshold, correlated with the global thermal noise level, to remove signal components that cannot be distinguished from i.i.d, zero-mean Gaussian data, which is attributable to thermal noise. Even though remaining signal components also contain some residual thermal noise (see discussion in Methods), the overall impact is a significant improvement in tSNR for NORDIC compared to Standard data (Figures 1C and Supplementary Figures 2 and 3) as well as to MPPCA (Supplementary Figure 16).

Difference of NORDIC *vs.* Standard images show only noise, which, when averaged over all the images in the fMRI times series demonstrates equivalence to the g-factor maps (Supplementary Figure 1), without evidence of edge effects or features of the imaged object; additionally, the FFT power spectra (Supplementary Figure 7) display only a broadband decrease in the magnitude of the spectrum without impacting the various peaks detected at specific frequencies associated with the stimulus presentation or physiologic fluctuations. These observations are consistent with the expectation that NORDIC suppresses random noise associated with the thermal noise of the MR measurement without perturbing the image.

T-values are a useful metric in evaluating functional mapping studies. Denoising algorithms inherently alter the dimensionality of the data and, consequently, the DFs of GLM computations. GLM’s DFs are crucial in computing p-values, though the correct computation of DFs for an fMRI time series is debated^43^. Here we do not attempt to address this issue, which is beyond the scope of this work as it relates not only to denoised time-series but is intrinsic to fMRI in general. We chose instead to compute our t-values using equation 2 (Methods) to provide a measure of activation relative to GLM residual noise. Thus, our activation maps based on t-rather than p-value thresholds, although we give the equivalent p value as a reference for the Standard reconstruction.

At the same t-threshold, the extent of voxels showing stimulus-invoked signal changes that pass the t-threshold, is considerably larger for the NORDIC processed single run (Figures 2 and 3; Supplementary Figures 4, 8-12) and equivalent to activation maps produced by concatenating 3 to 5 runs of the Standard data. This was also consistently observed for 3T and 7T data obtained with different resolutions, paradigms, and cortical regions (Supplemental Figures 8-19, 12). These observations are expected given the fact that NORDIC improves more than 2-fold the trial-to-trial precision of single-voxel PSC estimates while not impacting the magnitude of the PSC (Figures 3 and 4). Thus, NORDIC better estimates the stimulus evoked responses and does so in shorter runs in fMRI studies. Single trial responses represent a challenging SNR starved scenario and capturing them accurately with low single-trial variance is a highly desirable, yet seldomly achievable feat, especially in submillimeter resolution fMRI.

One of the most important features of NORDIC is its ability to preserve spatial precision of the individual images of the fMRI time series, as well as the precision of the functional response. Thermal noise associated with the MR process can and often is suppressed with spatial filtering, which smooths (i.e. blurs) the images, increasing the SNR and consequently the tSNR^44^; this improves the t-values (Supplementary Figures 8 and 11) and also, when applied with a Gaussian kernel, serves the purpose of making more valid the assumption of smoothness for FWER control based on random field theory (RFT) approaches widely used in the fMRI community. However, the resultant spatial blurring leads to an undesirable loss of spatial precision. NORDIC, on the other hand, suppresses thermal noise and has the same impact on t-values as spatial-smoothing (Supplementary Figures 8 and 11) but *without* spatial blurring of either the individual images themselves (Figures 5C, 6, and Supplementary Figure 14 and also see discussion in Methods) or the functional PSF estimates in the visual cortex (Figure 5A and 5B), yielding PSF values consistent with previous reports^22, 45^.

NORDIC can be said to improve the spatial specificity to neuronal activity changes by reducing false positives, and negatives. However, there could be additional benefits in specificity due to the ultrahigh resolutions enabled by NORDIC. At sufficiently high enough resolutions, the draining vein confound (e.g. see^46^) of GE BOLD fMRI is less of a problem because partial voluming and spatial averaging will be less and there would exist many voxels unaffected by this confound providing access to tissue responses, just like in optical imaging with intrinsic signals where blood vessels are visible but the high resolution permits visualization of the tissue responses in between blood vessels. There is, however, an additional advantage that can arise from the small voxel sizes achievable with NORDIC. GE BOLD fMRI is based on the voxel-wise measurement of signal *amplitude* after it is allowed to decay for an echo time TE with the rate constant 1/T_2_*. T_2_* strongly depends on intravoxel *B_0_* inhomogeneities, hence on neuronal activity because of the extravascular *B_0_* gradients generated by deoxyhemoglobin containing blood vessels. However, in the limit voxel dimensions become small compared to the spatial scale of these extravascular *B_0_* gradients, intravoxel inhomogeneities, hence their contribution to 1/T_2_*, also become small, reducing the detectability of stimulus/task induced alterations as an *amplitude* change in GE BOLD fMRI. For a given blood vessel, this limit is determined approximately by (δd/r_b_) where δd is the voxel dimension and r_b_ is the blood vessel radius, and the distance from the blood vessel since the gradient becomes rapidly shallower with increasing distance from the blood vessel. Thus, high resolutions enabled by NORDIC and other advances will lead to an intrinsic shrinkage of the spatial extent of the draining vein confound in GE BOLD fMRI and ultimately its suppression. Extravascular *B_0_* gradients still exist, of course. However, in this limit, they will show up as a phase difference among the different voxels. Such phase effects mixed with amplitude changes were already reported and used to account for large draining confound in GE BOLD fMRI^47^. As the resolutions increase, however, the amplitude effects will become smaller, leaving behind ultimately only the phase perturbation. Intravascular BOLD effect will still persist and will be a source of unwanted BOLD signals at lower magnetic fields like 3T^48^ but not at ≥7T where the very short T_2_ of blood assures its elimination^49^.

In the mammalian cortex there exist elementary cortical units of operation, consisting of several hundreds or thousands of neurons, that are spatially clustered and repeated numerous times in each cortical area. These mesoscopic scale ensembles are the focus of extensive research carried out in animal models by invasive techniques, such as optical imaging or electrophysiology. However, these techniques cannot be used in human studies because of their invasive nature. Therefore, the ability to generate functional maps at the level of these elementary units by MR methods is critically important and has been shown to be feasible^7-9^. However, current achievable resolutions (∼0.8 mm isotropic or non-isotropic voxels of equal volume) and the responses detected at such high resolutions are at best marginal. Overcoming this barrier with reasonable acquisition times has not been possible. For example, it has been possible to detect axis of motion features in the human MT^50^ but not the *direction of motion* subclusters that distinguish the motion in the two different directions along a given axis. It has been possible to demonstrate layer specific activations aimed at studying laminar organization but not with sufficient resolution to even distinguish three layers across the cortex without partial voluming and with Nyquist sampling; at least three (ideally more) distinct layers are required in order to clearly differentiate feedforward inputs arriving primarily into layer 4, local computations and cortico-cortical inputs shaping responses in layers 2/3, and outputs to other brain areas from layers 2/3, and 5/6.

The afore-described barrier and its limitations on neuroscientific research were recognized in the first report of the BRAIN Initiative Working group^6, 51^, which challenged the MR community to overcome it and achieve whole brain imaging studies with at least 0.1 µL voxel volumes (e.g. 0.46 or ∼0.5 mm isotropic resolution). We demonstrate here that this goal is achieved with NORDIC (Figure 6) at 7 Tesla and likely will soon be surpassed when multiplicative gains will be attained combining NORDIC with additional independent gains from acquisition methods, higher magnetic fields^52^, high channel count RF coils employed synergistically with very high magnetic fields^53, 54^ and image reconstructions methods (e.g.^55, 56^).

In this paper, we demonstrate an fMRI denoising approach to remove thermal noise inherent in the MR detection process, and markedly improve some of the most fundamental metrics of functional activation detection while crucially preserving spatial and functional precision. We demonstrate its efficacy for 7T mapping at high spatial resolution, as well as for 3T and 7T fMRI studies using the more commonly employed supra-millimeter spatial resolutions targeting different cortical regions activated by different stimuli and tasks. Importantly, as it specifically acts on Gaussian distributed noise, NORDIC is *complementary* as well as *beneficial* to denoising algorithms that primarily focus on structured, non-white noise removal. The cumulative gains are expected to bring in transformative improvements in fMRI, permitting higher resolutions at 3T, 7T and higher magnetic fields, more precise quantification of functional responses, faster acquisitions rates, significantly shorter scan times, and the ability to reach finer scale mesoscopic organizations that have been unreachable to date.

## Methods

### Image Reconstruction

*2D slice selective accelerated acquisitions*: For 2D acquisition with phase-encoding undersampling and/or simultaneous multislice (SMS)/Multiband (MB) acquisition, the GRAPPA and slice-GRAPPA reconstructions were used as outlined in^57^. A single kernel G^ch^_j_ is constructed for SMS/MB with/without phase-encoding undersampling such that for each slice, *j*, and channel, *ch*,

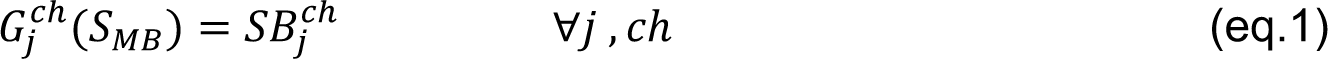

where S_MB_ denotes the acquired SMS/MB k-space, and SB^ch^_j_ denotes the reconstructed k-space for the slice *j* and channel *ch*. The kernels G^ch^_j_ are calculated similarly as in unbiased slice-GRAPPA from the measured individual slices SB_i_ with 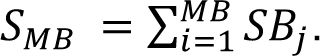

### 3D accelerated acquisitions

For 3D acquisitions with phase-encoding undersampling only, a gradient recalled echo (GRE) based Nyquist-sampled auto-calibration signal (ACS) reference acquired without slice-phase-encoding (a single slice-phase-encoding plane) was used. A Fourier transform was first applied along the slice-phase-encoding, and then k-space interpolation along the phase-encoding direction was performed with GRAPPA-weight calculated from the ACS reference.

### g-factor noise for image-reconstruction

g-factors were calculated building on the approach outlined in ^58^ for g-factor quantification in GRAPPA reconstructions and detailed in ^57^. The same ESPIRIT sensitivity profiles used for image reconstructions were also used for the determination of the quantitative g-factor.

### NORDIC PCA

Let 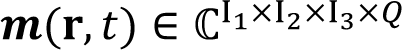 denote a complex-valued volumetric fMRI image series following an accelerated parallel imaging acquisition, where *Q* is the number of temporal samples and I_1_, I_2_, I_3_ the matrix size of the volume. The flow chart in Figure 7, adapted from^40^, illustrate the principles of NORDIC denoising of this dataset **m(r, t)** and the details of the noise model, locally low rank model, threshold selection, and patch averaging.

**Figure 7:**
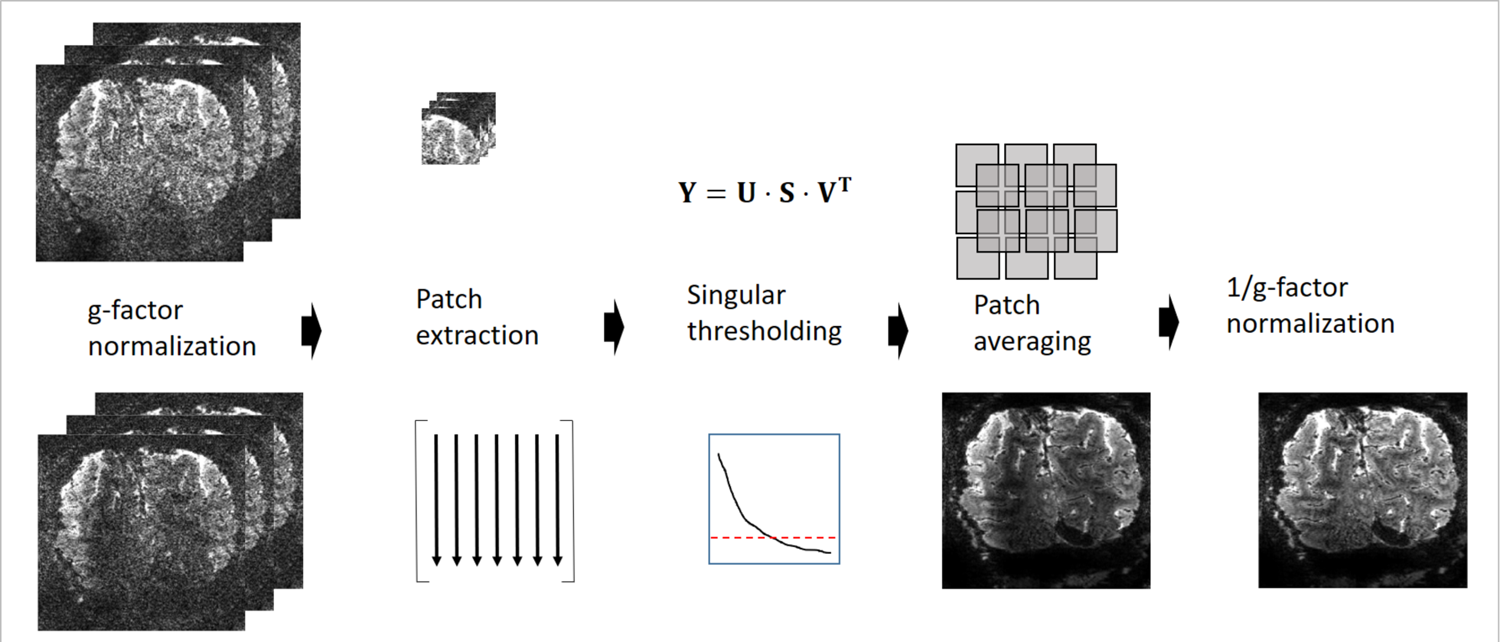
Flowchart of the **NORDIC** algorithm for a series m(r, τ). First, to ensure i.i.d. noise the series is normalized with the calculated g-factor kernels as m(r, τ)/g(r). From the normalized series, the Casorati matrix Y = [y_1_, ⋯, y_j_, ⋯, y_N_] is formed, where y_j_ is a column vector that contains the voxel values in each patch. The low-rank estimate of Y is calculated as Y_L_ = U ⋅ S_λthr_ ⋅ V^T^, where the singular values in S, λ(i) are set to 0 if λ(i) < λ_thr_. After re-forming the series m^LLR^(r, τ) with patch averaging, the normalization with the calculated g-factor is reversed as m^NORDIC^(r, τ) = m^LLR^(r, τ) ⋅ g(r).

### Noise Model

Images in MRI are inherently complex-valued but constructed as real-valued by using the magnitude of the images. This transformation changes the thermal Gaussian i.i.d. noise in the original measurement to be Rician for magnitude of coil-combined images or non-central Chi^2^ distributed when combining multiple magnitude images from different coils. Furthermore with parallel imaging reconstruction, the noise undergoes a spatially varying amplification, which is characterized by the geometry-factor, g(r). In NORDIC, a signal and noise scaling is performed on the complex valued data as m(r, t)/g(r) to ensure zero-mean and spatially identical noise in a given patch (Left-most column, Fig. 7). For NORDIC processing, a sensitivity weighted channel combination^59^ is applied to the accelerated dataset^57^ to maintain complex-valued Gaussian noise^60^ of the combined image, and the images are transformed to magnitude images only after denoising.

## Locally low rank Model

For locally low rank (LLR) processing, a fixed k_1_ × k_2_ × k_3_ patch is extracted from each volume in the series, and the voxels in each patch from each volume is vectorized as y_t_, to construct a Casorati matrix Y = [y_1_, ⋯, y_t_, ⋯, y_Q_] ∈ ℂ^MxQ^ with M = k_1_ × k_2_ × k_3_, and Q representing the number of volumes (time points) in the fMRI time series. The concept of NORDIC is to estimate the underlying matric X in the model where Y = X + N, and N ∈ ℂ^MxQ^, where N is additive Gaussian noise.

LLR modelling assumes that the underlying data matrix X has a low-rank representation. For NORDIC, k_1_ × k_2_ × k_3_ is selected to be a sufficiently small patch size so that no two voxels within the patch are unaliased from the same acquired data for the given acceleration rate^40^, ensuring that the noise in the pixels of the patch are all independent. LLR methods typically implement the low-rank representation by singular value thresholding (SVT). In SVT, singular value decomposition is performed on Y as U ⋅ S ⋅ V^H^, where the entries of the diagonal matrix S are the ordered singular values, λ(j), j ∈ {1, …, N}. Then the singular values below a threshold λ(j) < λ_thr_ are changed to λ(j)=0 while the other singular values are unaffected. Using this new diagonal matrix S_λthr_, the low-rank estimate of Y is given as Y_L_ = U ⋅ S_λthr_ ⋅ V^H^.

## Hyperparameter Selection

While the threshold in NORDIC is chosen automatically without any empirical tuning, the method itself has hyperparameters related to the patch size that determine the size of Casorati matrices. In NORDIC, k_1_ × k_2_ × k_3_ is selected with M ≈ 11 ⋅ Q, and k_1_ = k_2_ = k_3_, as determined heuristically in Moeller et. al.^40^. We note that the choice of patch size with a M: Q ratio of 11:1, can be more challenging to accommodate for long fMRI runs since Q, is the number of samples in the time series, especially in light of the requirement that no two voxels within a patch are unaliased from the same acquired data. For whole brain rsfMRI, as in the Human Connectome Project, for example, M ≈ 11 ⋅ Q can be maintained. If there is an issue fulfilling this requirement, the geometry of the patch may be adjusted to something different than k_1_ = k_2_ = k_3_.

The patches can be either 2D or 3D, and while 2D patches may better fit with the temporal dynamics of the acquisition, the data independence constraint of no two voxels within the patch being unaliased from the same acquired data can be challenging. For longer series, the constraint of M ≈ 11 ⋅ Q may either in itself not be satisfied simultaneously with the data independence, or it may be further difficult in the presence of phase-encoding ghosting e.g. from fat or eddy currents. 3D patches are less limited in this regard and also better capture spatially similar signals.

## Noise Model and Threshold Selection

The distribution of the singular values of a random noise matrix N is well-understood if its entries are i.i.d. zero-mean. The threshold that ensures the removal of components that are indistinguishable from Gaussian noise is the largest singular value of the noise matrix N. While this threshold is asymptotically specified through the Marchenko-Pastur distribution, for practical finite matrix sizes, we numerically estimate this value via a Monte-Carlo simulation^40^. To this end, random matrices of size *M*×*Q* are generated with i.i.d. zero-mean entries, whose variance match the experimentally measured thermal noise, σ^2^, in **Y**. The thermal noise level can be determined after g-factor normalization from an appended acquisition without RF excitation (a noise acquisition) to the series, or from a region of interest outside the brain devoid of signal contributions or it can be determined from a receiver noise pre-whitening acquisition. In this paper the first method of utilizing an additional noise-acquisition has been employed. Then the empirical mean value of the largest singular value is used as the numerical threshold.

## The Degree of Noise Removal

Though NORDIC removes zero-mean, i.i.d. Gaussian noise, it does not remove all of it. This can be explained more formally by considering one of the *M x Q* Casorati matrices we are trying to denoise based on the model **Y** = **X** + **N**. According to our model, **Y** is the observed noisy data, **N** is a matrix whose entries are zero-mean, i.i.d. Gaussian, and **X** is the low-rank data matrix. More concretely, the low-rank condition states rank(**X**) = *r* << min{*M*, *Q*} = *Q* (latter equality due to our choice of *M*). For ease of explanation, also assume that all non-zero singular values of **X** are sufficiently above the noise level. Thus, when the singular value decomposition of **Y** is performed, it will have *r* singular values that contain a combination of signal component from **X** and noise component from **N**, while the remaining (*Q – r)* singular values will only have contributions from noise **N**. Since the thresholding is performed at the level of the largest singular value of the noise matrix, NORDIC will remove the noise from all these (*Q – r*) noise components, as they cannot be distinguished from zero-mean i.i.d Gaussian noise (i.e. random noise). On the other hand, the *r* singular values that are above the threshold will be unaffected by NORDIC processing. However, these *r* singular values have contributions from both noise and signal components, though these components will be dominated by the signal. Thus, the final denoised estimate generated from these singular values and their corresponding singular vectors will have residual Gaussian noise in them. Since *r* << *Q* due to low-rank assumption, majority of the thermal noise is removed by virtue of thresholding (*Q* – *r)* singular values, but a small amount of thermal noise that are on the remaining *r* singular components will remain in the final estimate. As a side note, this remaining thermal noise will be further reduced due to patch averaging in processing, but this effect is difficult to quantify.

## Patch averaging

The patches arising from these thresholded Casorati matrices are combined by averaging^61^ overlapping patches to generate the denoised image series m^LLR^(r, τ). The averaging of patches can be performed with patches having different geometries, i.e. k_1_, k_2_, k_3_, and the averaging can be identically weighted or weighted by the number of non-zero λ’s. In NORDIC for fMRI, direct averaging with identical weights is used, similar to the previous use of NORDIC in dMRI, where it was shown that there was no difference from using weighted averaging^40^. The patch-averaging is itself a denoising step^62^ which reduces the residual contributions of noise. In NORDIC for fMRI, with typically Q > 100, and M > 1000, we used patch averaging with 25-50% overlap, and the difference between this and using all combinations of patches was minimal, but led to substantial savings in computational time.

Finally, to obtain m^NORDIC^(r, t) the denoised volumes m^LLR^(r, t) are multiplied back with the g-factor map g(r) to correct the signal intensities.

## Spatial Blurring

It may seem counter-intuitive that noise can be removed without introducing spatial blurring. The main idea behind the locally low-rank decomposition is to separate out the noisy Casorati matrix **Y** into two components as **Y** = **X** + **N**, where **X** is assumed to be low-rank, and **N** is Gaussian noise. Then the algorithm thresholds to remove all principal components of **Y**, whose singular values are below the threshold that is automatically determined in NORDIC by the noise level. This will remove both contributions from **N** and from **X**. This is analogous to the concept of image compression, where part of the data is removed (e.g. some of the DCT coefficients in JPEG compression), but the end result is visually indistinguishable from the uncompressed image, as long as the compression level is not too high. In this analogy, the compression is done via removing some of the components of the low-rank **X**, but due to its low-rank property, this does not fundamentally alter its visualization. Additionally, the compression level in conventional image compression is analogous to the SNR/threshold level in our method. A numerical simulation of the threshold and patch size relative to zero-mean Gaussian noise was performed in Moeller et al^40^

### Participants

To test the impact of NORDIC on fMRI, we acquired 10 data sets on four (2 females) healthy right-handed subjects (age range: 27-33), with different stimulation paradigms, acquisition parameters and field strengths (see *Stimuli and Procedure* and *MRI Imaging Acquisition and Processing* paragraphs). All subjects had normal, or corrected vision and provided written informed consent. The study complied with all relevant ethical regulations for work with human participants. The local IRB at the University of Minnesota approved the experiments.

### Stimuli and procedure

We tested the impact of NORDIC on fMRI across 4 experimental paradigms:

1. Block design visual stimulation
2. Fast event related visual stimulation design
3. Fast event related auditory stimulation design
4. Resting state.

#### 1. Block design visual stimulation

We implemented standard block design visual stimulation paradigms (see Figure 1A) for 4 acquisition types. These included the two 3T fMRI studies, the 0.8 mm isotropic resolution 7T fMRI and the 7T 0.5 mm isotropic resolution fMRI datasets (see *MRI Imaging Acquisition and Processing* paragraph). The experimental procedure consisted of a standard 12 s on, 12 s off for the 7T 0.8 mm isotropic voxel acquisitions, and for the 3T datasets, and a 24 s on, 24 s off for the 7T 0.5 mm isotropic voxel acquisitions (see Figure 1A). The difference in block length between the 2 resolutions was implemented to account the difference in volume acquisition time between the 0.8 mm iso (i.e. volume acquisition time = 1350 ms) and the 0.5 mm iso acquisitions (i.e. volume acquisition time = 3652 ms). The stimuli consisted of a center (i.e. target) and a surround square checkerboard counterphase flickering (at 6 Hz) gratings (Figure 1A) subtending approximately 6.5 degrees of visual angle. Stimuli were centered on a background of average luminance (25.4 cd/m^2^, 23.5-30.1). Stimuli were presented on a Cambridge Research Systems BOLDscreen 32 LCD monitor positioned at the head of the 7T scanner bed (resolution 1920, 1080 at 120 Hz; viewing distance ∼89.5 cm.) using Mac Pro computer. Stimulus presentation was controlled using Psychophysics Toolbox (3.0.15) based codes. Participants viewed the images through a mirror placed in the head coil.

Each run lasted just over two and a half minutes for the 0.8 mm 7T and the 3T acquisitions (i.e. 118 volumes at 1350 ms TR) and just over 5 minutes for the 0.5 mm 7T acquisitions (85 volumes at 3654 ms volume acquisition time), beginning and ending with a 12 s or 24 s red fixation dot centered on a gray background. Within each run, each visual condition, target and surround, was presented 3 times. For the 0.5 mm iso data sets, we collected 8 experimental runs; for the 0.8 mm iso 7T and the two 3T data sets, participants underwent 8 runs, 2 of which were used to compute the region of interest and excluded from subsequent analyses. Participants were instructed to minimize movement and keep fixation locked on the center fixation dot throughout the experimental runs. For the 0.8 mm 7T acquisition on S3, run 8 had to be discarded due to excessive movement.

#### 2. Fast event related visual design

The visual fast event related design consisted 6 runs of a face detection task, with a 2 s on, 2 s off acquisition. Each run lasted approximately 3 min and 22 s and began and ended with a 12 s fixation period. Importantly, we introduced 10% blank trials (i.e. 4 s of fixation period) randomly interspersed amongst the images, effectively jittering the ISI. Stimulus presentation was pseudorandomized across runs, with the only constraint being the non-occurrence of 2 consecutive presentations of the same phase coherence level.

Behavioral metrics, including reaction time and responses to face stimuli indicating participants’ perceptual judgments (i.e. face or no face) were also recorded.

We used grayscale images of faces (20 male and 20 female). We manipulated the phase coherence of each face, from 0% to 40% in steps of 10%, resulting in 200 images (5 visual conditions x 20 identities x 2 genders). We equated the amplitude spectrum across all images. Stimuli approximately subtended 9 degrees of visual angle. Faces were cropped to remove external features by centering an elliptical window with uniform gray background to the original images. The y diameter of the ellipse spanned the full vertical extent of the face stimuli and the x diameter spanned 80% of the horizontal extent. Before applying the elliptical window to all face images, we smoothed the edge of the ellipse by convolving with an average filter (constructed using the “fspecial” function with “average” option in MATLAB. This procedure was implemented to prevent participants from performing edge detection, rather than the face detection task, by reacting to the easily identifiable presence of hard edges in the face images.

#### 3. Fast event related auditory design

Stimuli consisted of sequences consisting of four tones. For each sequence, tones were presented for 100 ms with a 400 ms gap in between them (sequence duration 1.6 s). The sequences were presented concomitantly with the scanner noise (i.e. no silent gap for sound presentation was used) and 36 tone sequences were presented in each run, a session consisted of 10 runs of about 6 minutes each. Tone sequences were presented following a slow-event related design with an average interval of 6 TR’s (ranging between 5 and 7 TR’s, TR = 1.6 s).”

#### 4. Resting state

The resting state acquisition consisted of four 10 minute runs. Data were obtained at 3T with 3T HCP acquisition parameters (see section below). No stimulus presentation occurred and participants were instructed to stay still, minimize movements and fixate on a visible crosshair.

## MR Imaging Acquisition and Processing

### 7T Acquisition parameters

All 7T functional MRI data were collected with a 7T Siemens Magnetom System with a single transmit and 32-channel receive NOVA head coil.

We collected 4 variants of T2*-weighted images with different acquisition parameters, tailored to the different experimental needs. Specifically, for *block design* visual stimulus paradigm at 7T we collected 0.5 mm iso voxel (T2*-weighted 3D GE EPI, single slab, 40 slices, TR 83 ms, Volume Acquisition Time 3654ms, 3-fold in-plane undersampling along the phase encode direction, 6/8ths in plane Partial Fourier, 0.5 mm isotropic nominal resolution, TE 32.4ms, Flip Angle 13°, Bandwidth 820Hz). The 0.8 mm iso voxel acquisition used T2*-weighted 2D GE SMS/MB EPI, 40 slices, TR 1350 ms, Multiband factor 2, 3-fold in-plane undersampling along the phase encode direction, 6/8ths Partial Fourier, 0.8mm isotropic nominal resolution, TE 26.4ms, flip Angle 58°, Bandwidth 1190Hz. For the *auditory event related design*, we used a comparable submillimeter acquisition protocol (2D GE SMS/MB EPI 42 slices, TR 1600 ms, Multiband factor 2, 3-fold in-plane undersampling along the phase encode direction, 6/8ths Partial Fourier, 0.8 mm isotropic nominal resolution, TE 26.4 ms, Flip Angle 61°, Bandwidth 1190Hz)

For the *visual fast event related design*, we used the 7T HCP acquisition protocol (2D GE SMS/MB EPI, 85 slices TR 1s, Multiband factor 5, 2-fold in-plane undersampling along the phase encode direction, 7/8ths Partial Fourier, 1.6 mm isotropic nominal resolution, TE 22.2 ms, Flip Angle 51°, Bandwidth 1923Hz)

## 3T Acquisition parameters

We recorded data employed the *block design visual stimulus* paradigm using 2 sequences varying in resolution: Acquisition sequence 1 used the 3T HCP protocol parameters (72 slices, TR= 0.8s, Multiband= 8,no in-plane undersampling 2mm isotropic, TE =37ms, Flip Angle= 52°, Bandwidth =2290 Hz/pixel). Acquisition sequence 2 parameters were 100 slices, TR= 2.1s, Multiband= 4, in-plane undersampling factor = 2, 7/8 Partial Fourier, 1.2mm isotropic, TE= 32.6ms, Flip Angle= 78°, Bandwidth= 1595Hz/pixel

For the resting state data we used the acquisition sequence 1 detailed above (i.e. the 3T HCP protocol).

For all acquisitions, flip angles were optimized to maximize the signal across the brain for the given TR. For each participant, shimming to improve B0 homogeneity over occipital regions was conducted manually.

T1-weighted anatomical images were obtained on a 3T Siemens Magnetom Prisma^fit^ system using an MPRAGE sequence (192 slices; TR, 1900 ms; FOV, 256 x 256 mm; flip angle 9°; TE, 2.52 ms; 0.8 mm isotropic voxels). Anatomical images were used for visualization purposes and to define the cortical grey matter ribbon. This was done in BrainVoyager via automatic segmentation based on T1 intensity values and subsequent manual corrections. All analyses were subsequently confined within the gray matter.

## Functional data Preprocessing

All 7T Functional data preprocessing was performed in BrainVoyager. Preprocessing was kept at a minimum and constant across reconstructions. Specifically, we performed slice scan timing corrections for the 2D data (sinc interpolation), 3D rigid body motion correction (sinc interpolation), where all volumes for all runs were motion corrected relative to the first volume of the first run acquired, and low drift removals (i.e. temporal high pass filtering) using a GLM approach with a design matrix continuing up to the 2^nd^ order discrete cosine transform basis set. No spatial nor temporal smoothing was applied. Functional data were aligned to anatomical data with manual adjustments and iterative optimizations.

3T dicom files were converted using dcm2niix^63^. All subsequent 3T functional data preprocessing was performed in AFNI version 19.2.10^64^. Conventional processing steps were used, including despiking, slice timing correction, motion correction, and alignment to each participant’s anatomical image.

EPI data were aligned to T1 weighted images. For all multiband data sets (i.e. all acquisitions other than the 3D 0.5 mm iso images), anatomical alignment was performed on the Single Band Reference (SBRef) image which was acquired to calibrate coil sensitively profiles prior to the multiband acquisition and has no slice acceleration or T1-saturation, yielding higher contrast^65^.

## GLMs and tSNR

Stimulus-evoked functional maps were computed in BrainVoyager for all 7T datasets and in AFNI for the 3T datasets. ROI definition and contrast maps were also computed using these software. Subsequent analyses (i.e. ROI based and functional point spread function measurements) were performed in MatLab using a set of tools developed inhouse.

Temporal tSNR was computed by dividing the mean (over time) of the detrended time-courses by its standard deviation independently per voxel, run and subject.

To quantify the extent of stimulus evoked activation, we performed general linear model (GLM) estimation (with ordinary least squares minimization). Design matrices (DMs) were generated by convolution of a double gamma function with a “boxcar” function (representing onset and offset of the stimuli). We computed both single trial as well as condition-based GLMs. The latter, where DMs had one predictor per condition, were used to assess the differences in extent

and magnitude in activation between NORDIC and Standard images. The former, where the DMs had one predictor per trial per condition, produced single trials activation estimates that were used to assess the stability (see “*NORDIC vs. Standard statistical analyses”* paragraph below) of the responses evoked by the target condition for each voxel within the left retinotopic representation of the target in V1 (see below).

## ROI definition

Out of the 8 recorded runs, 2 runs (identical for each reconstruction type) were used to define a region of interest (ROI). Specifically, we performed a classic GLM on 4 concatenated runs (2 reconstructed with NORDIC and 2 with the standard algorithm) and computed the differential map by contrasting the t-values elicited by the target to that elicited by the surround. While this approach may overinflate statistical power and misrepresent the size of the ROI, it also ensures identical ROIs across reconstructions, which was the main goal in this case. GLM t-values can be thought of as beta estimates divided by GLM standard error according to this equation:

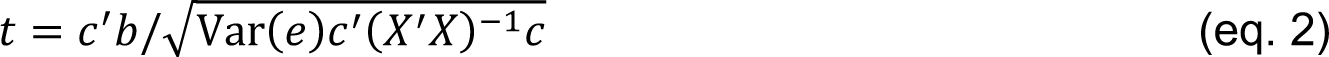

where b represents the beta weights, c is a vector of 1, −1 and 0 indicating the conditions to be contrasted, e is the GLM residuals and X the design matrix. We then thresholded this map (p<.05 Bonferroni corrected) to define the left hemisphere retinotopic representation of the target stimulus within the grey matter boundaries. This procedure was implemented to provide an identical ROI across reconstruction types, however, it resulted in effectively doubling the number of data points available, which could not be treated as independent anymore. To partially account for this, we adjusted the GLM degrees of freedom used to compute the t-maps to be equal to those of 2 rather than 4 runs.

## GLMs for experimental runs

Independently per reconstruction type, for the condition-based scenario, GLMs were performed for each single run as well as for multiple runs (i.e. concatenating 2 or more experimental runs and design matrices to estimate BOLD responses). For the multiple run scenarios, we estimated the percent signal change beta weights and the related t-values for 2, 3, 4, 5 and 6 runs. For each *n*-run GLM, we computed independent GLMs for all possible run combinations (see the “*Comparing extent of activation*” paragraphs for more details).

## NORDIC vs. Standard statistical analyses

In order to evaluate the impact of NORDIC denoising on BOLD based GE-EPI fMRI images, the following analyses were performed. Standard tSNR was computed as described earlier. To assess statistically significant differences in average tSNR across reconstruction types, we first computed the mean tSNR (using the 20% trimmed mean, which is more robust to extreme values^23^) across all voxels in the brain for each of the 8 runs. We then carried out 2-tailed paired sample t-tests between average tSNRs for NORDIC and Standard images across all runs.

Moreover, to test for statistically significant differences in stimulus-evoked BOLD amplitudes and noise levels across reconstruction algorithms, we compared the ROI voxel mean percent signal change beta estimates and related t-values elicited by the target condition independently per subject. We used the 18 responses elicited by the 3 stimulus presentations within each of the 6 runs. To account for the fact that trials within each run are not independent, while the runs are, we implemented a Linear Mixed-Effect Model in Matlab (The Mathworks Inc, 2014) according to the equation:

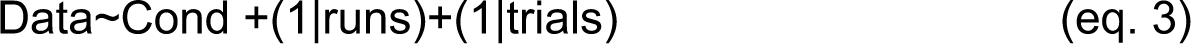

Linear Mixed-Effect Model allows estimating fixed and random effects, thus allowing modeling variance dependencies within terms. Model coefficients were estimated by means of maximum likelihood estimation.

To assess differences in the precision of BOLD PSC estimates across reconstructions, we computed the cross-validates R2 for single runs GLMs. This was achieved by deriving the beta weights using a given “training” run, and testing how well these estimates predicted single voxel activation for all other “test” runs. Single voxel cross validated R2 (also known as coefficient of determination) was computed according to the equation:

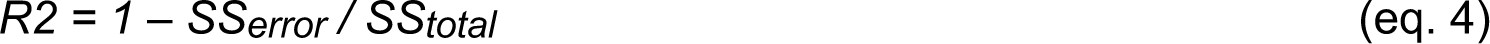

and, in our specific case

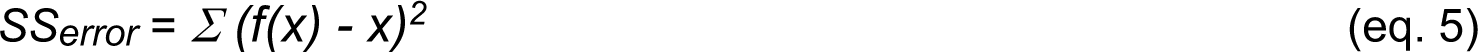

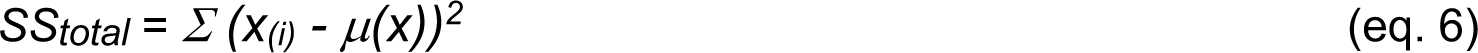

In eq. 5 and 6, *x* is the empirical time-course of the test run, *x(i)* represents the i^th^ point of the empirical time-course of the test run *x*, *μ(x)* is the time-course mean, and *f(x)* is the predicted time course computed by multiplying the design matrix of a given training run by the beta estimates derived on a different test run.

We computed all possible unique combinations of training the model on a given run and testing on all remaining runs, leading to 15 R2s per voxel. To infer statistical significance, we carried out paired sample t-tests across the 15 cross-validated R2 (averaged across all ROI voxels) for NORDIC and Standard images.

To assess the stability and thus the reliability of single trials response estimates we computed the standard deviation across percent signal change amplitudes elicited by each single presentation (i.e. single trial) of the target stimulus for every run, voxel and reconstruction type. To infer statistical significance between these stability estimates for NORDIC and Standard, independently per subject we carried out 2 tests: 1) we performed 2-tailed paired sample t-tests across runs; 2) we computed 95% bootstrap confidence intervals as follows. First, for a given subject, we computed the difference between the single trials’ standard deviations of NORDIC and Standard data. For each bootstrap iteration, we then sampled with replacement the runs, computed the mean across the sampled runs and stored the value. We repeated this operation 10000 times, leading to 10000 means. We sorted these 10000 means and selected the 97.5 and the 2.5 percentiles (representing the 95% bootstrapped confidence intervals of the difference). Statistical significance was inferred when 95% bootstrap confidence interval did not overlap with 0.

## Comparing extent of activation

We further compared the extent of activation across reconstructions for the GLMs computed on 1 and multiple runs by quantifying the number of active voxels at a fixed t-value threshold. To this end, we computed the t-map for the contrast target > surround. For each GLM, we then counted the number of significant voxels at t ≥ |5.7| (corresponding to p<.05 Bonferroni corrected for the Standard images) within the ROI. As we intended to understand and quantify the difference in extent of activation between NORDIC and Standard reconstructions, we compared GLM computed on 1 NORDIC run versus 1, 2, 3, 4, 5 and 6 runs of Standard GLMs. To ensure that any potential difference was not related to run-to-run variance, we implemented the following procedure. Firstly, we computed GLMs for all possible unique run combinations. This led to six data points for single run GLMs (i.e. 1 GLM per experimental run), 15 data points for GLMs computed on 2 concatenated runs (e.g. runs 1-2; 1-3;1-4;1-5;1-6; 2-3 etc.), 20 data points for GLMs computed on 3 concatenated runs; 15 data points for GLMs computed on 4 concatenated runs; 6 data points for GLMs computed on 5 concatenated runs and 1 data point for the GLM computed on 6 concatenated runs. For each run combination, we counted the significant number of active voxels at our statistical threshold and stored those numbers. Within each n-run GLM (where n represents the number of concatenated runs), we then proceeded to compute 95% bootstrap confidence interval on the mean of the active number of voxels across all possible run combinations. This was achieved by sampling with replacement the number of significantly active voxels estimated for each combination of runs and computing the mean across the bootstrap sample. We repeated this operation 1000 times to construct a bootstrap distribution and derive 95% bootstrap confidence interval^23^. This procedure not only ensured sampling from all runs, but it also decreased the impact of extreme values^23^.

## Quantifying BOLD images Smoothness

Global smoothness estimates from each reconstruction prior to preprocessing (‘pre’) and following all data preprocessing, just prior to the GLM (‘post’). This was performed using 3dFWHMx from AFNI^64^ using the ‘-ACF’ command. The data were detrended using the default settings from 3dFWHMx with the ‘-detrend’ command. As we are interested in the smoothness within the brain, we also used the ‘-automask’ command in order to generate an intensity-based brain mask, based on the median value of each run. This method iterates through various background clipping parameters to generate a contiguous brain only volume, that excludes the external areas of low signal. The spatial autocorrelation is estimated from the data using a Gaussian plus mono-exponential model, which accounts for possible long-tail spatial autocorrelations found in fMRI data. This estimated FWHM, in mm, from this fitted autocorrelation function is used as an estimate of the smoothness of the data. This estimate was derived for all of the runs, excluding the held-out runs used for ROI creation. For each subject, smoothness was averaged within each stage across the 6 experimental runs to evaluate if global smoothness was markedly increased due to the reconstruction method. Paired sample t-tests were carried out between estimated FWHM parameters for the NORDIC and Standard reconstructions to infer statistical significance.

## Functional point spread function

Functional point spread function (PSF) was computed according to^22^. We estimated the BOLD functional PSF on all individual runs independently for the Standard and NORDIC reconstructions. In brief the analysis was implemented as follows: We first identified the anterior most retinotopic representation of the target’s edge in V1 separately in Standard and NORDIC reconstructed data. This was achieved by computing the contrast target > surround on all runs concatenated within each group (Standard vs. NORDIC) and identifying those voxels showing differential BOLD closest of 0 (Figure 5). Then, using BrainVoyager, we flattened this portion of the cortex to produce Laplace-based equipotential grid-lines in the middle of the cortical ribbon. To increase the precision of the PSF measurement, we upsampled the BOLD activation maps to 0.1 mm isotropic voxel. Independently per run, we then drew 10 traces orthogonal to the retinotopically anterior most edge of the target. We estimated the BOLD functional PSF on all individual runs independently for the Standard and NORDIC reconstructions. In brief the analysis was implemented as follows: We first identified the anterior most retinotopic representation of the target’s edge in V1 separately in Standard and NORDIC reconstructed data. This was achieved by computing the contrast target > surround on all runs concatenated within each group (Standard vs. NORDIC) and identifying those voxels showing differential BOLD closest of 0 (Figure 5). Then, using BrainVoyager, we flattened this portion of the cortex to produce Laplace-based equipotential grid-lines in the middle of the cortical ribbon. To increase the precision of the PSF measurement, we upsampled the BOLD activation maps to 0.1 mm isotropic voxel. Independently per run, we drew 10 traces orthogonal to the retinotopically anterior most edge of the target. We then superimposed these traces to the activity elicited by the target condition and, from the target’s edge, we measured the slope of BOLD amplitude decrease along the traces. PSF was quantified by fitting a model to the mean of the 10 traces consisting of a step-function (representing infinitely precise PSF) convolved with a gaussian^22^ with 3 free parameters. The 3 parameters were the width of the gaussian (representing functional precision – see^22^, the retinotopic location of the edge and a multiplicative constant. Parameter fitting was performed in Matlab using the *lsqcurvefit* function, with sum of squares as stress metric. Paired sample t-tests across the 8 runs were then carried out between the Gaussian widths for NORDIC and Standard images to infer statistical significance.

## Resting state analysis

We collected 4 sequential runs of resting state. Each run was 10 minutes in length, with the subject fixating on a crosshair throughout. Minimal processing steps, performed with AFNI, were applied to the Standard and NORDIC data. These included slice timing correction and motion correction to the first volume of the first run of the Standard data for both Standard and NORDIC data. For both reconstructions, motion was computed (and corrected) relative to the first volume of the first run of the Standard data. Next, we regressed out the 6 estimates of motion parameters and polynomials up to 5^th^ order. A spherical seed, with radius of 3mm was placed in the medial prefrontal cortex, corresponding to a location within the Default Mode Network. The extracted seed time course for each run was used to generate a map of Pearson’s r values, corresponding to the correlation of each voxel in the brain with the seed timeseries (i.e. seed-based correlation).

## Denoising algorithms comparison

We compared the performance of NORDIC to that of other denoising strategies on the 0.5 mm isotropic functional data, which, amongst the many datasets in this paper, represents the one most greatly affected by thermal noise and therefore an ideal candidate for NORDIC. Specifically, we evaluated the performances of a global PCA based algorithm^26^ (PCAwn), and a local PCA based algorithm DWIDenoise (DWIdn) ^27^, on both magnitude dicoms and complex dicoms. DWIdn is a publicly available implementation of the MPPCA method^27^. PCAwn was implemented following Thomas et al. ^26^ by first selecting all voxels in the brain by image intensity thresholding on the average series, and then applying a SVD on the Casorati matrix of the whole volume. Following the SVD, each of the left singular basis vectors was evaluated for signal contributions using the multi-taper analysis^25, 26^, and an empirical threshold, determined from the ratio of the power to the standard deviation of the power spectrum from the multi-taper analysis ^25, 26^, was utilized to select and remove components which only contributed to the thermal noise as in ^26^. DWIdn. is a local PCA method designed to select and remove components which only contributed to the thermal noise using an objective threshold derived from the Marchenco-Pastur distribution for random matrices. DWIdn was applied both on magnitude only and on complex dicoms using the MRtrix3 toolbox; http://www.mrtrix.org, with its default optimized settings^27^. To obtain complex dicoms, we converted phase images to radians and then combined them with magnitude images using mrcalc, a tool also part of MRtrix3. Of these methods NORDIC and DWIdn are the most similar, but presenting a number of importance difference, including, but not limited to, the approach of the threshold selection and the optimization of local patch-size (see^40^ for more details).

The impact of denoising methods on fMRI data entailed comparing a number of metrics all described in the previous pages. These included the t-maps for the contrast target > surround; the distribution of these t-values on an ROI hand drawn on the co-registered T1, approximately corresponding to the representation of the target region in V1; smoothness metric as implemented in AFNI and the impact of the different methods on single EPI image quality. The reason for hand drawing the ROI rather than deriving it from the maps themselves, was to ensure no bias towards a specific denoising algorithm. The ROI was further constrained to only include values within the brain for the EPI images, to account for potential misregistration across modalities. The results of these comparisons are presented in Supplementary Figure 16.

## ACKNOWLEDGEMENTS

The authors thank Prof. Kendrick Kay, University of Minnesota, for helpful discussion and comments. This work was supported by NIH grants U01EB025144 (K.U.), P41 EB027061 (K.U.), P30 NS076408 (K.U.) and RF1 MH116978 (E.Y.), and RF1 MH117015 (Geoffrey Ghose, University of Minnesota), and NSF grant CAREER CCF-1651825 (M.A.).

## AUTHOR CONTRIBUTIONS

LV, EY, KU designed the experiments. LV and LD acquired data. LD analyzed the 3T data; FdM analyzed the auditory data; LV carried out all remaining analyses. SM and MA conceptualized the denoising methodology, SM wrote the denoising algorithm and executed it on the acquired data. LV, KU, SM, MA wrote the paper. All authors discussed the results, the manuscript, and edited the manuscript.

## DATA AVAILIBILITY

The data that support the findings of this study are available from the corresponding author upon reasonable request, subject to human subjects IRB limitations. Source data are provided with this paper

## CODE AVAILIBILITY

The codes that support the findings of this study will be available here: https://github.com/SteenMoeller/NORDIC_Raw<u>; and on zenodo (</u>DOI:10.5281/zenodo.5032437)

## SUPPLEMENTARY MATERIAL

The supplemental information presented in this section falls into two categories**: I)** Supplementary Figs. 1 through 7, showing additional analyses and results based on the 0.8 mm isotropic resolution 7 Tesla data presented in the main body of the paper to supplement the conclusions reached from these 7T data; **II)** additional data sets and discussion demonstrating the wide applicability of NORDIC across field strengths, cortical regions, stimulation and/or task paradigms, and acquisition strategies.

### I. Supplementary Figures based on the 7T data presented in the main body of the paper

**Supplementary Figure 1.**
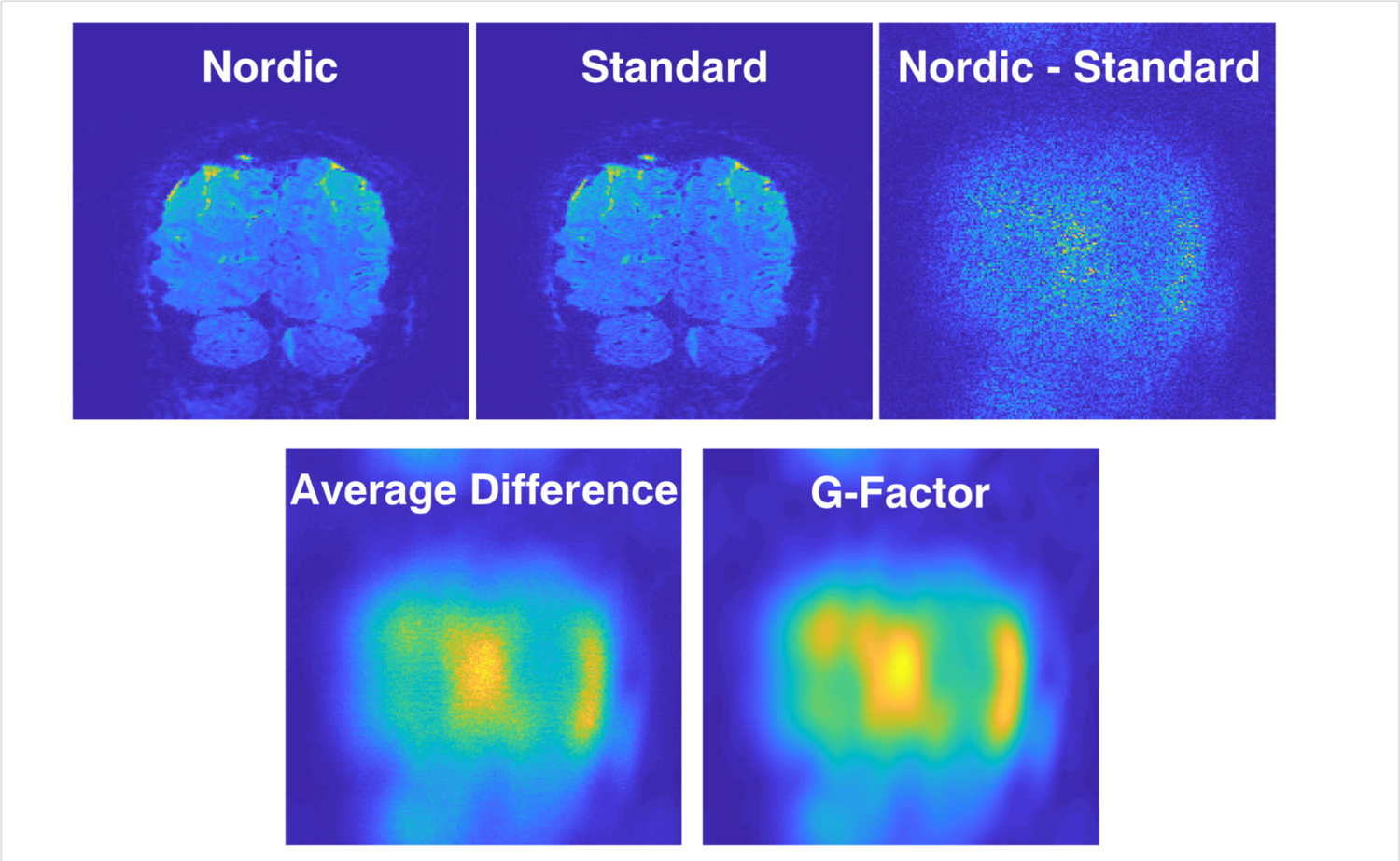
Difference images obtained by subtracting Standard reconstructed image from the same image reconstructed with NORDIC. The top row shows data from a single slice and a single time point in the fMRI time series obtained from subject 1 with 0.8 mm isotropic resolution voxels and the block design target and surround retinotopic stimuli presented in the main text (Figs. 1 through 5). NORDIC and Standard reconstructed images of this slice are shown together with a difference between these reconstructions (top row, right most panel). Note how the structured noise reflects the G-factor (shown in the rightmost panel second row), which, incidentally, is comparable to the average of Nordic-Standard across all volumes. The lower row, left panel shows the same NORDIC minus Standard difference for the same slice but now averaged for all time points in the fMRI time series. Lower row right panel shows the g-factor map. These results demonstrate that there are no structures (e.g. edges) in the difference maps reflecting the features of the images they come from; such edge effects would be expected if there is blurring of the image by NORDIC Furthermore, when averaged over all data in the fMRI time series, the resulting image (lower row, left panel) looks like the g-factor map (lower row, right panel), as would be expected if NORDIC is removing the instrumental thermal noise amplified by the g-factor.

**Supplementary Figure 2.**
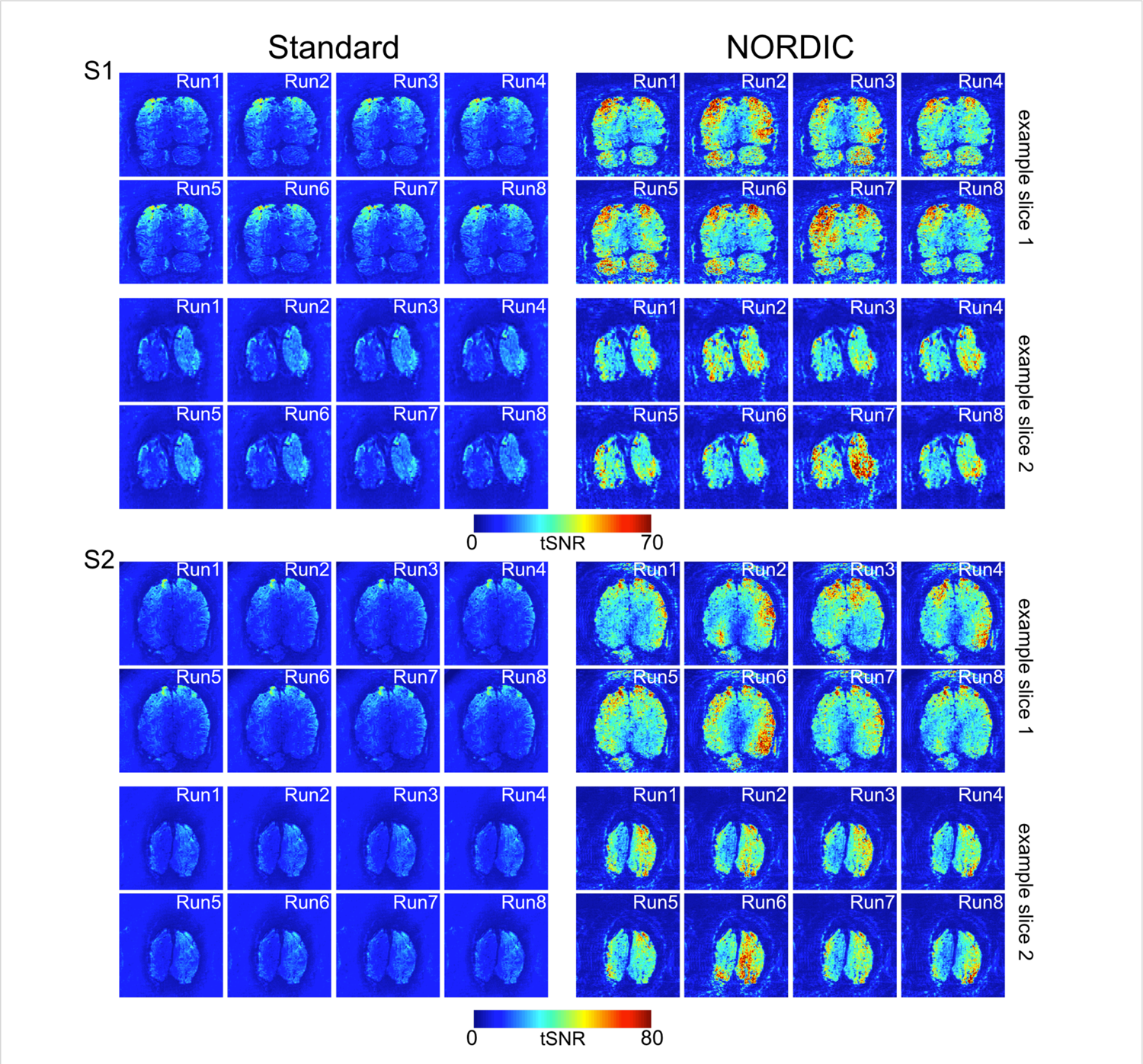
Single runs temporal signal-to-noise ratio (tSNR) maps. tSNR maps of 2 exemplar slices for two subjects for all 8 different runs for Standard (left) and NORDIC (right) reconstructions. Slice 1, represents one of the anterior most slices in the covered volume; slice 2 is an occipital slice that includes a portion of V1.

**Supplementary Figure 3.**
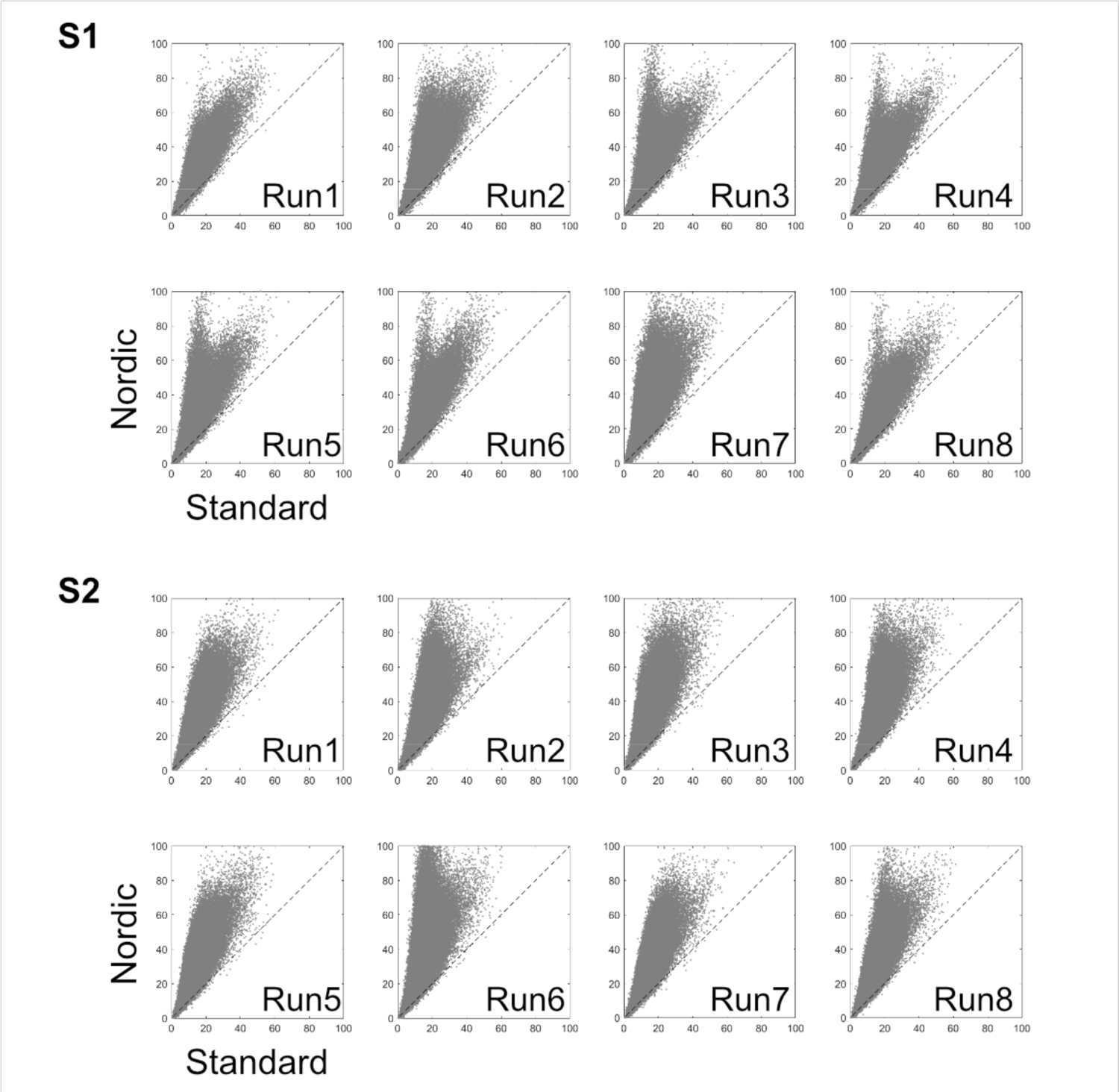
Single run temporal signal-to-noise ratio (tSNR) scatter plots. Single run scatterplots of the temporal SNR of NORDIC vs Standard for all brain voxels for 2 representative subjects (S1 and S2, in Figure 2, main manuscript).

**Supplementary Figure 4.**
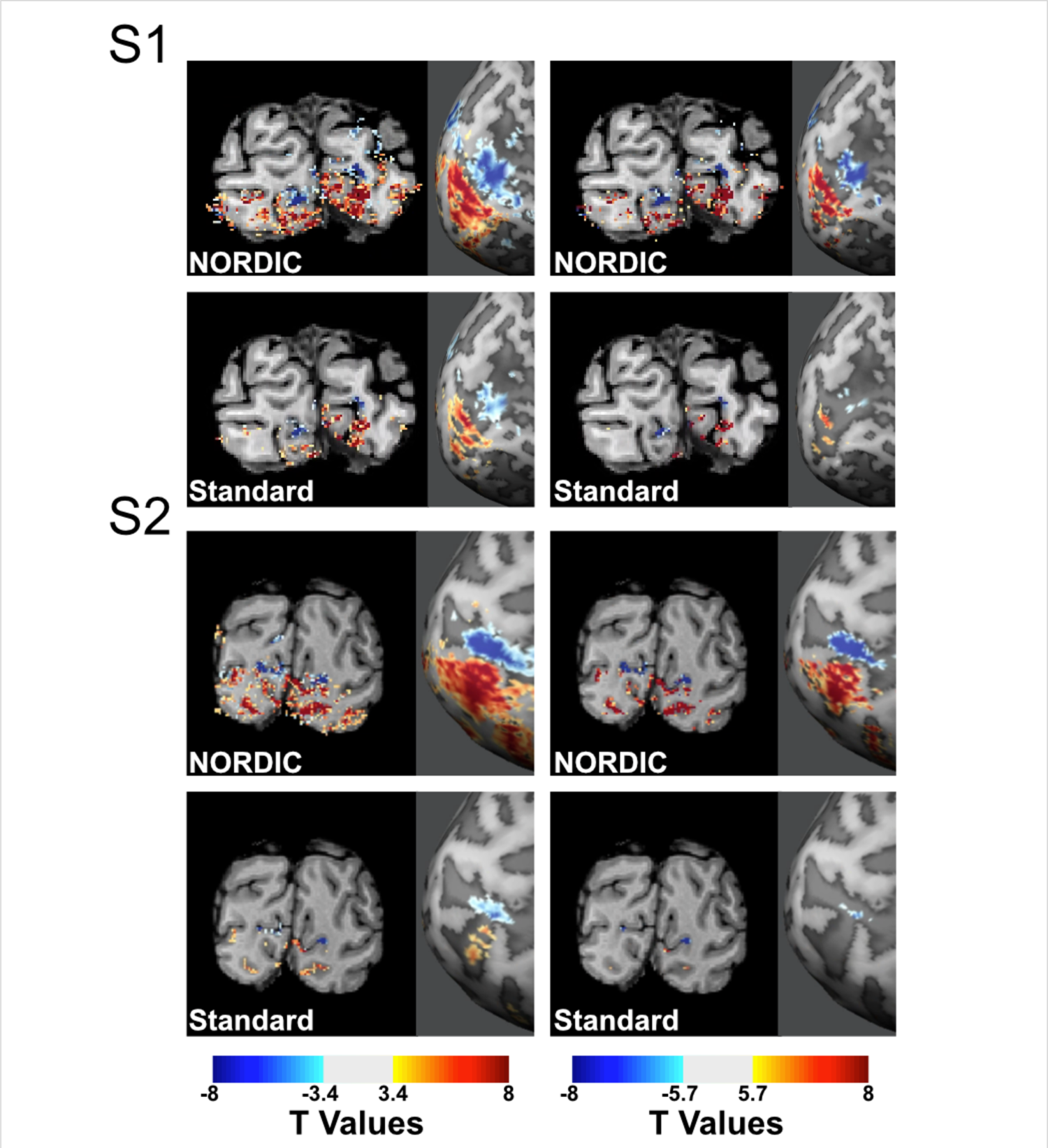
NORDIC vs. Standard t-maps. Examples of NORDIC and Standard reconstructed t-maps superimposed onto T1 weighted anatomical images for two subjects (S1 and S2, in Figure2, main manuscript) obtained from a single ∼2.5 min. For each subject, the images on the top row represent NORDIC reconstruction and the lower row the Standard reconstruction. The t-maps were computed by contrasting the activation elicited by the target (red) versus that elicited by the surround (blue) condition for a single representative run in volume space (left) and on inflated brains (right). The two different columns show different thresholds, specifically, t ≥ 3.4 (left column) and t ≥ 5.7 (right column); for the Standard reconstruction. For the Standard reconstruction. These t-values (from a 2-sided paired sample t-test) correspond to p<0.01 (uncorrected) and p < 0.05 (Bonferroni corrected).

**Supplementary Figure 5.**
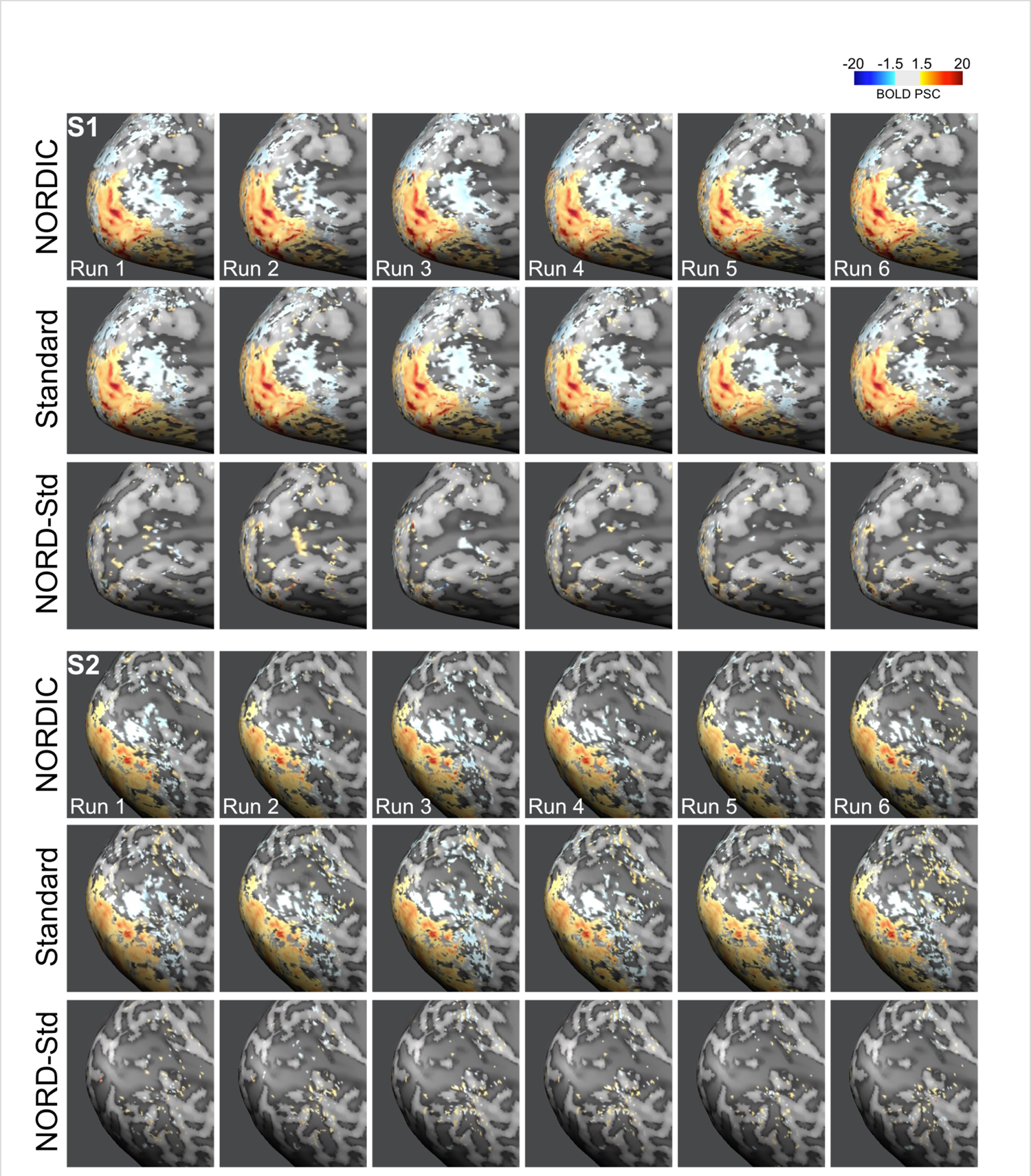
Single run percent BOLD signal change (PSC) maps for two exemplar subjects. For each subject (S1 and S2 in Figure 2, main manuscript) the top row portrays the NORDIC single run BOLD PSC maps elicited by the target condition. The second row is equivalent to the first, but for Standard images. The bottom row shows the BOLD difference between NORDIC and Standard. As evident in this figure, BOLD PSC maps are highly comparable across reconstructions.

**Supplementary Figure 6.**
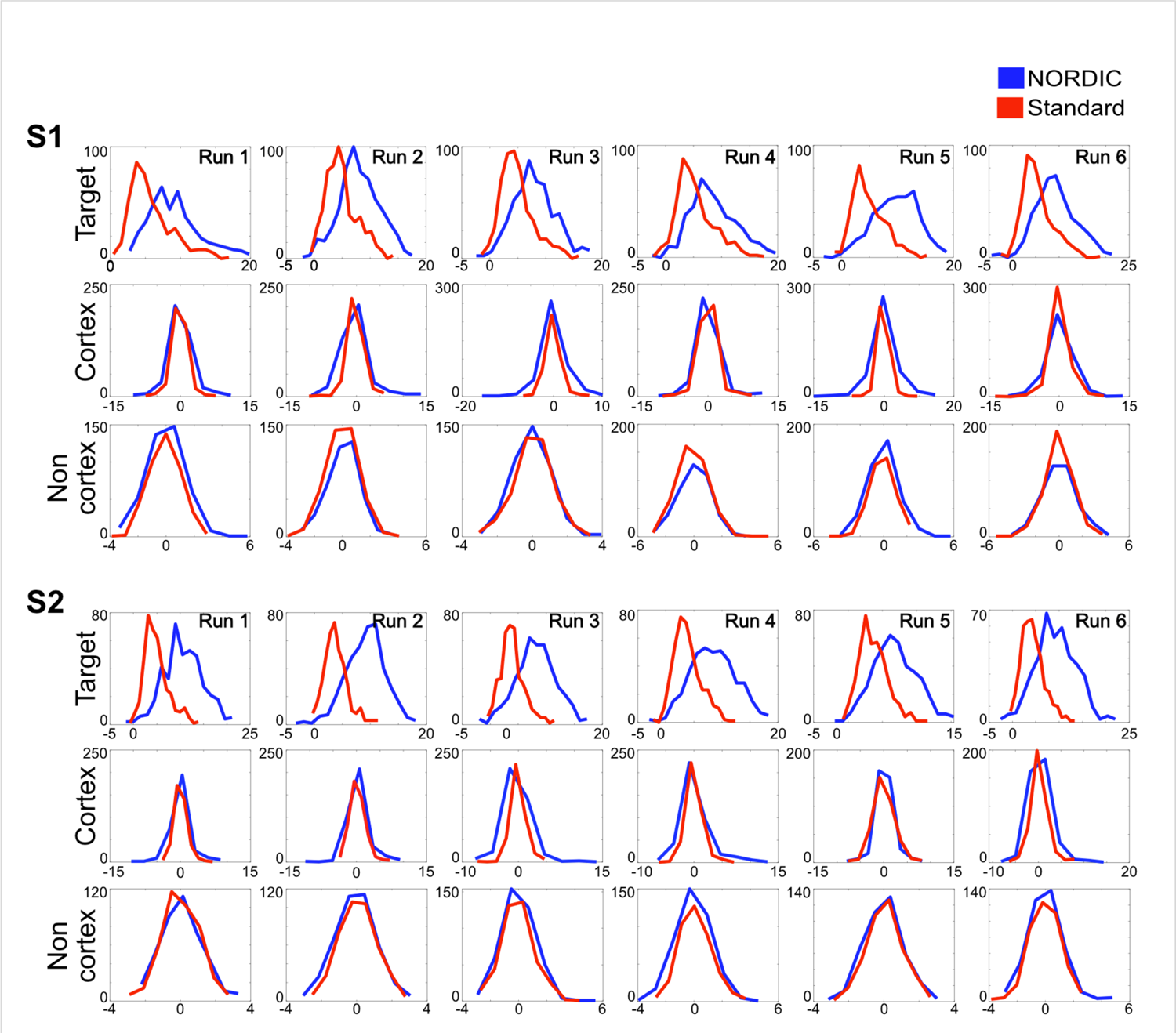
NORDIC vs. Standard t-values distributions. NORDIC (blue) and Standard (red) t-value distribution for the contrast Target > 0 (see methods) for 3 ROIs. In all plots, the x axis depicts the magnitude of the t-values, while the y axis shows the voxel count. For each subject (S1 and S2 in Figure 2 main manuscript) the top row shows the histogram of the t-values for all the voxels within the target ROI. The second row shows the histogram of the t-values for a number of randomly selected voxels within the cortex not belonging to the target ROI. The third row shows the histogram of the t-values for a number of randomly selected voxels outside the cortex. For the cortex and non-cortex ROIs, the number of voxels was chosen to match that of the target ROI. The t-values distribution for NORDIC and Standard are highly non-overlapping for the target voxels. In light of the comparable PSC amplitudes across reconstructions, this set of figures indicates a substantial noise reduction for NORDIC images. The histograms for voxels within the cortex but outside the target ROI or the non-cortical voxels are essentially overlapping, demonstrating that they are not perturbed by NORDIC processing, as should be the case.

**Supplementary Figure 7.**
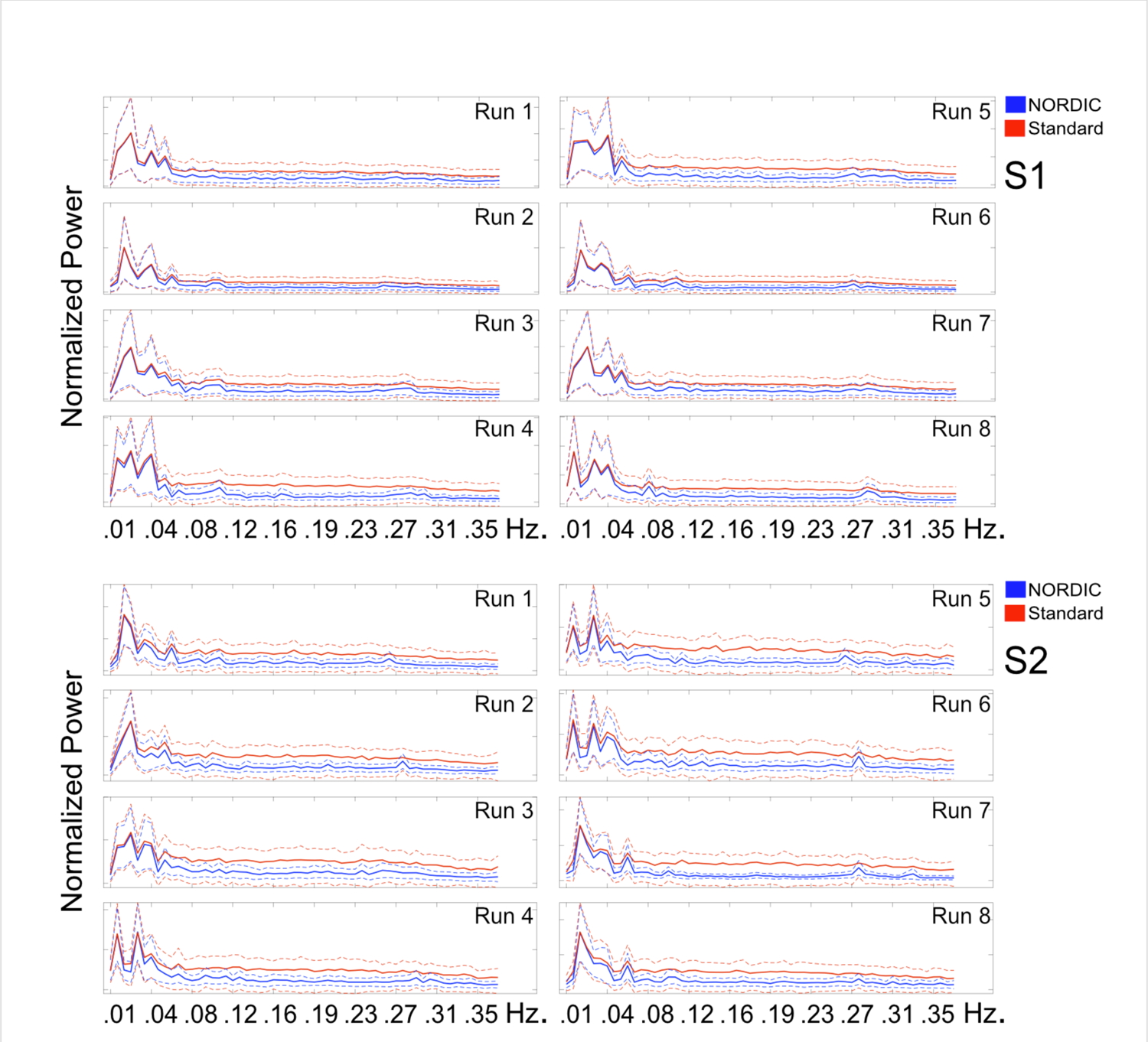
NORDIC vs. Standard Fast Fourier Transform (FFT). Supplementary Figure 7 shows FFT for all runs of 2 exemplar subjects (S1 and S2 in Figure 2 of the main manuscript) for NORDIC (blue) and Standard (red) images (dotted lines show the standard deviation across voxels). FFT was computed independently per voxel within the target ROI. The magnitude output of the FFTs were then averaged across all target voxels to produce the above power spectra. The x axis portrays frequencies in Hertz. For NORDIC (but not Standard) power spectra, a peak at approximately at 0.27 Hz – representing the respiratory frequency - is visible for both subjects and most runs. These figures confirm that NORDIC denoising operates in the thermal noise regime, not impacting the peaks in the FFT spectrum but causing a general frequency independent reduction in amplitude, consistent with suppressing white noise. They also illustrate that structured physiological noise, such as respiration, becomes more easily detectable after NORDIC, as expected. Lower frequencies, representative of neuronal responses, remain largely unaffected, indicating that NORDIC denoising does not compromise neural BOLD activity.

### **II.** Demonstrating applicability of NORDIC, across field strengths, spatial resolutions, cortical regions, stimulation and/or task paradigms, and acquisition strategies

#### ***i)*** NORDIC applications at 3 Tesla

In the manuscript, we presented only high resolution 7 Tesla fMRI data. However, vast majority of the functional imaging studies are carried out at 3 Tesla using supra-millimeter resolutions. Therefore, one of the most important indicators for the wide applicability of the NORDIC method would be to demonstrate that it can substantially improves such 3 Tesla data depending on the details of the acquisition protocol. Here, we demonstrate significant gains NORDIC imparts on the 3 Tesla data acquired with the Human Connectome Project (HCP) ^1, 2^ protocol (Supplementary Fig. 8) and a modification of the HCP protocol to achieve higher resolution with a lower MB factor and longer TR (Supplementary Figure 9). The stimulation paradigm was same retinotopically arranged target and surround visual stimulation paradigm described in the main manuscript (see Fig. 1A in the Manuscript).

**Supplementary Figure 8:**
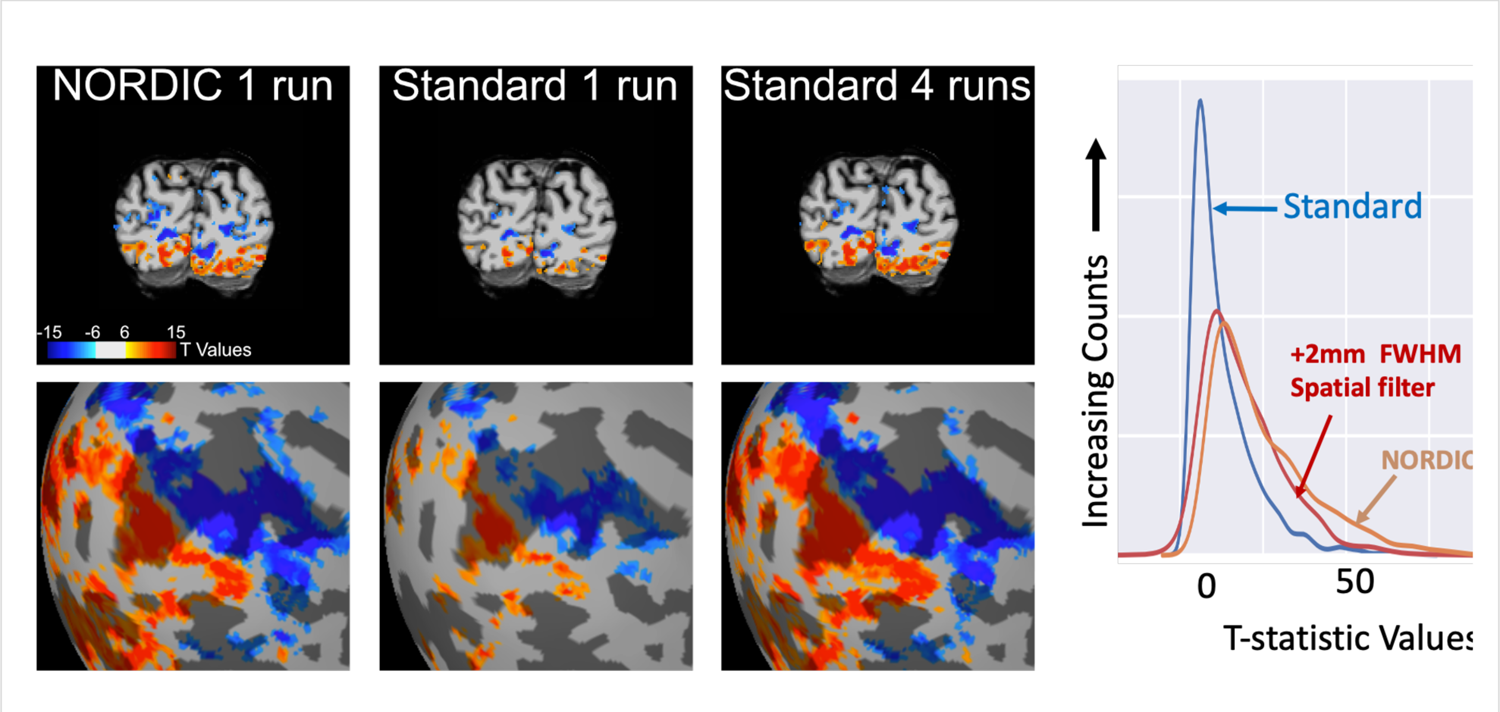
The effect of NORDIC denoising on an fMRI study carried out at 3 Tesla using the standard acquisition parameters of HCP: Spatial resolution= 2 mm isotropic; TR= 800 ms; Simultaneous Multislice(SMS)/Multiband (MB) EPI acquisition with MB factor of 8. Stimulus employed was identical to that implemented in the main study described in the manuscript (see Methods), where flickering gratings for target or surround were presented in 12 second blocks interleaved by 12 second fixation periods for ∼2.5 min for each run. The figure shows t-maps with t value ≥|6| for the contrast target (red) > surround (blue) in volume space for a representative slice in the visual cortex (top row), and in the inflated cortical surface (bottom row). T values >|15| appear as the same color as t=|15|. Approximately four runs with standard reconstruction (∼10 min of data) are required to achieve the extent of activation comparable a single NORDIC run (∼ 2.5 min of data). The right-most panel shows the t-value distribution for NORDIC, Standard reconstruction, and Standard reconstruction with 2 mm Full Width Half Maximum (FWHM) spatial smoothing, within a region of interest (ROI) created using the largest cluster from 8 runs of the data with Standard reconstruction and threshold of t=3.297, for the target *vs*. surround contrast. This ROI contained 2762 voxels. This figure demonstrates the benefit of NORDIC denoising with supra-millimeter acquisition protocols at 3 Tesla, further highlighting the wide applicability and relevance of NORDIC denoising.

As shown in Supplementary Fig. 8, the use of 4 concatenated runs representing ∼10 mins of data are needed to generate an fMRI map (t-statistics map with thresholding) equivalent to that obtained from a single run (∼ 2.5 min of data) using NORDIC denoising. The functional maps obtained with NORDIC using a single run is virtually identical on a pixel-by-pixel basis to that obtained with Standard 4 runs.

The t-statistics are also displayed in Supplementary Fig. 8 (right-most panel) for NORDIC and spatially smoothing with a 2 mm Full Width Half Maximum (FWHM) filter. The impact of this spatial smoothing filter on the t-statistics is similar, though not as good as to NORDIC. Unlike spatial smoothing, however, NORDIC achieves this improvement without degradation of spatial resolution (data presented in the main manuscript; also see discussion in Supplementary Fig. 11).

Looking into the future, one of the major impacts of the NORDIC approach will likely be to enable 3 Tesla studies at higher spatial resolutions relative to what has been achievable to date. We demonstrate this potential in Supplementary Fig. 9, using 1.2 mm isotropic resolution, lower MB factor 4, which may be preferred at 3T with a 32 channel coil, which results in a longer and more conventional TR.

**Supplementary Figure 9:**
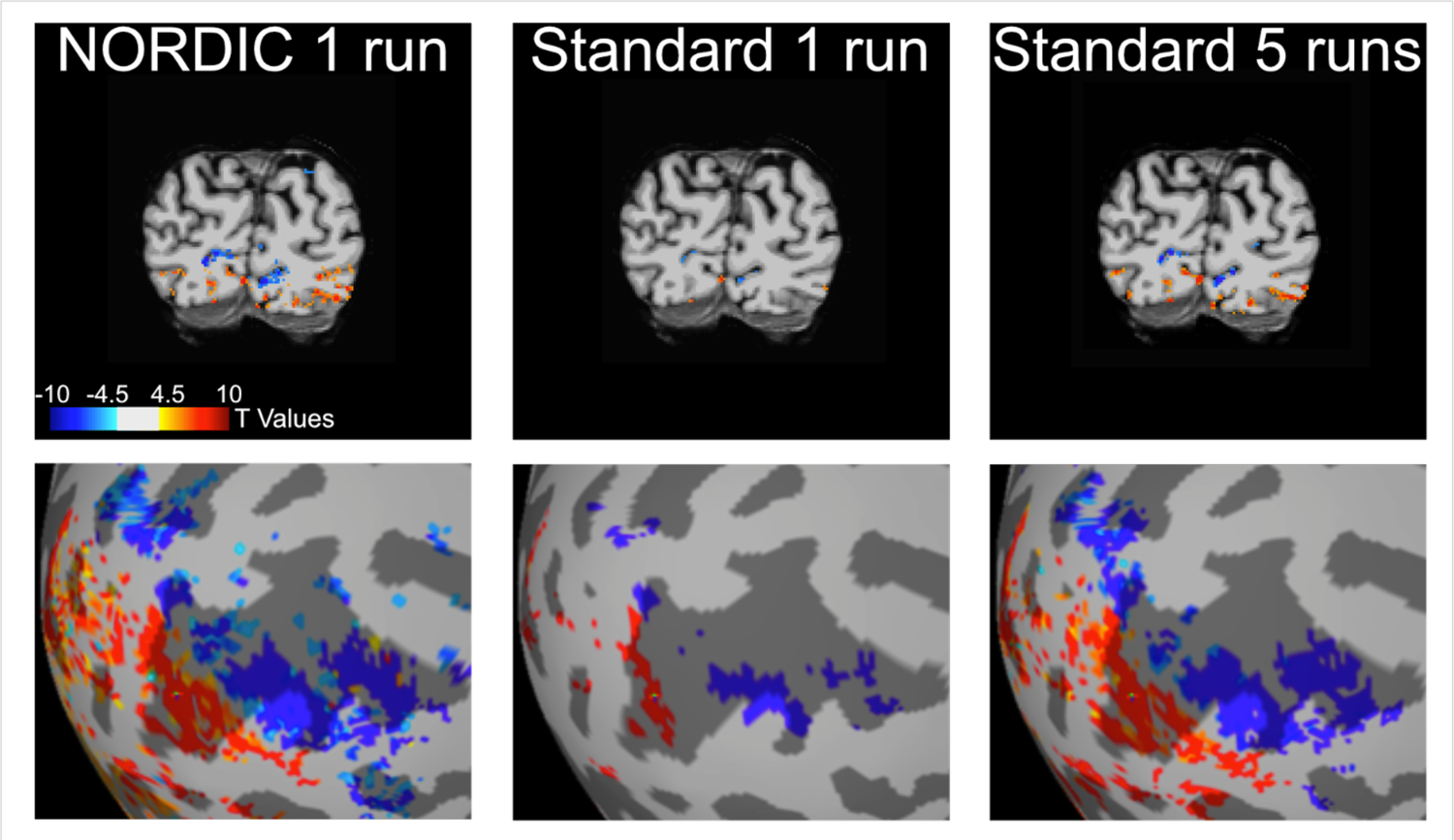
The effect of NORDIC denoising on an fMRI study carried out at <u>3 Tesla</u> using 1.2 mm isotropic resolution (TR 2100 ms; MB factor= 4; iPAT 2). The figure show t-maps with a t-threshold of ≥ |4.5| for the contrast target (red) > surround (blue). Because the higher resolution lengthens the echotrain length with deleterious contributions to image distortions and signal drop out, this acquisition employed parallel imaging along the phase encode direction (iPAT=2), and reduced the parallel imaging along the slice direction by using MB=4 as opposed MB=8 of the standard HCP protocol (Supplementary Fig.8). The stimulus and the presentation paradigm were the same as in Supplementary Fig.8 and identical to that implemented in the main study (see Methods), where flickering gratings for target or surround were presented for 12 seconds blocks interleaved by 12 seconds fixation periods. The figure shows t-maps for the contrast target (red) > surround (blue) in volume space, for a representative slice in the visual cortex (top row) and in inflated cortical space (bottom row). Approximately 5 Standard runs (∼12 minutes of data) are required to achieve the extent of activation comparable to a single NORDIC run (∼2.5 minutes of data). Given the significant differences in acquisition parameters relative to the HCP 3T data (Supplementary Fig.8), these results demonstrate the benefits of NORDIC in highly different MR context, further advocating for the generalizability of the technique.

The challenge of this higher resolution at 3 Tesla becomes evident with the functional map obtained from a single run processed with the Standard approach (Supplementary Fig. 9, middle column); this functional map is missing most of the activated territory evident in the data shown in Supplementary Fig. 8. However, the large activated territories seen in Supplementary Fig. 8 are largely recovered in NORDIC reconstruction of a single run (Supplementary Fig. 9, left column) or with Standard reconstruction after concatenating five runs, representing a much longer (∼12 min) data acquisition (Supplementary Fig. 9, right most column).

#### ***ii)*** EVENT RELATED Designs with CONVENTIONAL (Supra-millimeter) and HIGH (sub-millimeter) SPATIAL RESOLUTIONS and using complex COGNITIVE tasks (Face detection and gender discrimination)

In order to demonstrate the gains achieved by NORDIC are applicable at 7T data with i) more standard (i.e. ≥ 1 mm isotropic) spatial resolutions, ii) different stimuli or tasks, iii) different paradigms, and even iv) different regions of the cortex, we turned again to the HCP acquisition protocols. The HCP had a 7 Tesla component^1, 2^, and this component employed 1.6 mm isotropic resolution, TR=1 s, SMS/MB acquisition with MB=5, iPAT (undersampling in phase encoded direction) = 2. We used these acquisition parameters at 7T with a face detection paradigm where participants viewed degraded images of faces (ranging from 0% to 40% image phase coherence in steps of 10% increments) while performing a face detection task; the visual stimulation paradigm employed was a *fast event related* design where every image was presented for 2 seconds, followed by a 2 second fixation period (with 10% blank trials). The activation maps portrayed in Supplementary Fig. 10 show the T value maps with a t-threshold of t ≥ |4| for the mean activation to all 5 visual conditions. In this case the cortical region shown is the face fusiform area in the inferior temporal lobe.

**Supplementary Figure 10:**
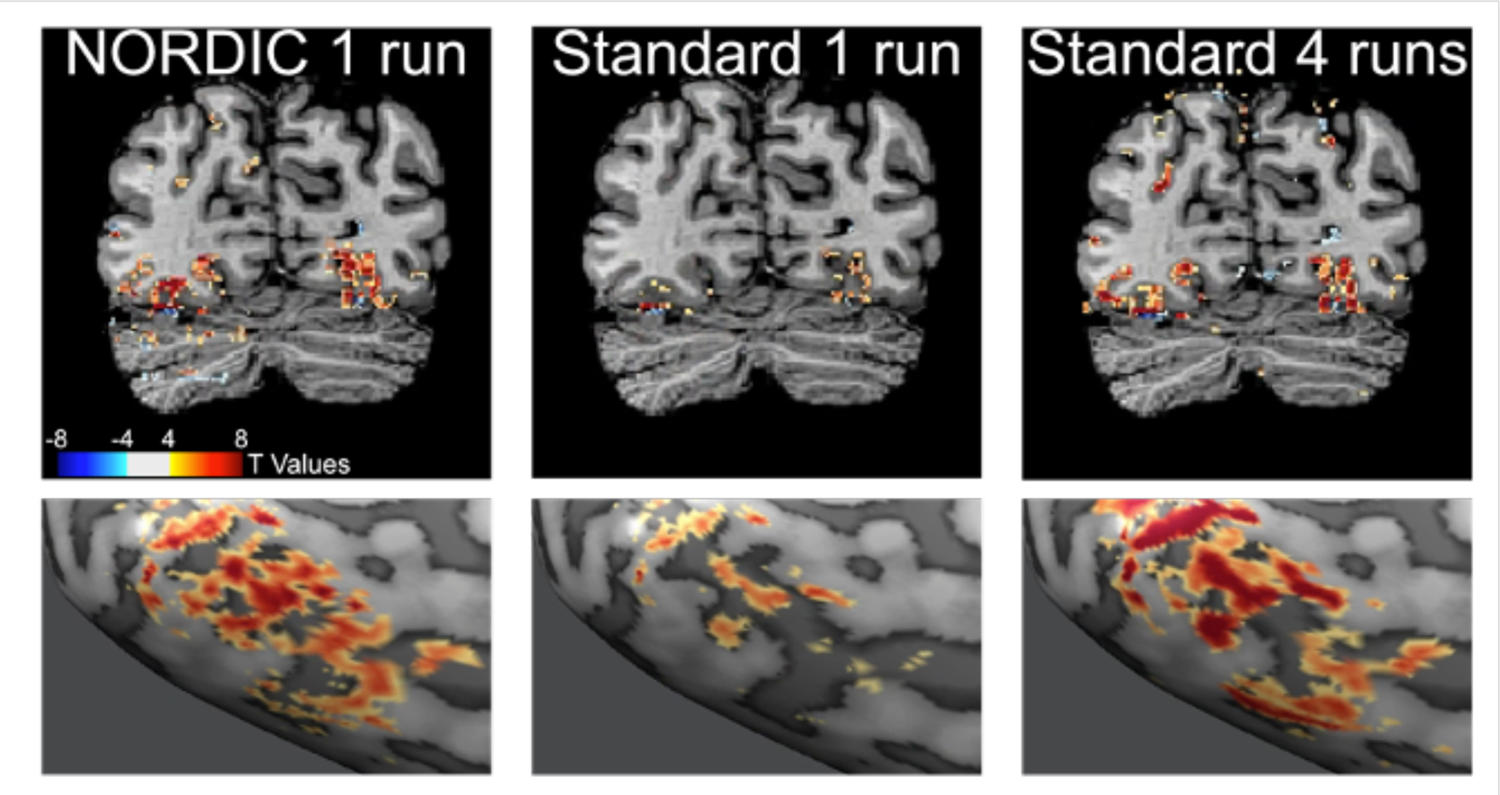
The impact of NORDIC denoising on 7T data acquired with a fast event related design and standard HCP 7T fMRI protocol (i.e. 1.6 mm iso voxels; TR 1000 ms; MB 5; iPAT 2). The event related paradigm used 2 seconds stimulus-on, 2 seconds stimulus-off. A 12 s stimulus-off baseline period was also utilized at the beginning and at the end of the rapid, 2 second on-off alternating epochs. During the stimulus on periods, the participants viewed degraded images of faces (i.e. ranging from 0% to 40% image phase coherence in steps of 10% increments) while performing a face detection task. The figure shows t-maps (t ≥ |4|) for the mean activation of all conditions vs. the mean of all baseline periods when nothing was shown; the top row shows this activation map in volume space for a single slice and the lower row displays it on the inflated cortical surface in the fusiform face area. Four Standard runs (∼13 minutes of data) are required to achieve the extent of activation comparable to a single NORDIC run (∼3 minutes and 20 seconds of data). The HCP acquisition protocol implemented here represents a gold standard for supra-millimeter 7T fMRI; therefore, these results highlight the wide-ranging advantages conferred by NORDIC denoising across acquisition and experimental protocols.

Again, we see that single run processed with NORDIC provides a functional map that is virtually equivalent to the map obtained using multiple runs (in this case 4) concatenated and processed with Standard reconstruction algorithms, and both are far superior to what is seen with a single run processed with Standard reconstruction (Supplementary Fig.10).

Supplementary Figure. 11 compares the effect of NORDIC on the t-statistics against the effects of spatial filtering (smoothing) for an *event related paradigm* similar to that shown in Supplementary Fig. 10, but acquired with higher spatial resolution (0.8 mm isotropic) using face presentations during a gender discrimination task. Supplementary Fig.11 illustrates that applying a 0.8 mm FWHM spatial smoothing on the raw data improves the t-statistics significantly, as in the case of the 3T data shown in Supplementary Fig. 5, albeit with the effective spatial resolution degraded by approximately a factor of two in this case. NORDIC, on the other hand, performs comparable to or slightly better than spatial filtering with respect to improvements in the t-statistics (Supplementary Fig. 8), but does not degrade spatial resolution (see data presented in the main body, and also Supplementary Fig. 16).

**Supplementary Figure 11:**
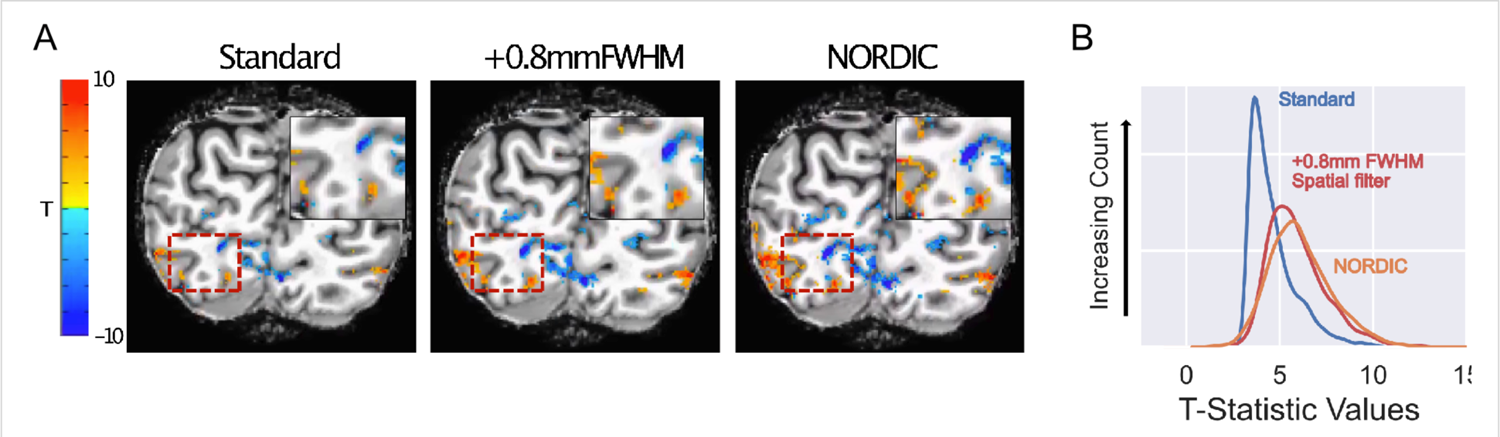
T-map (panel A) and their distributions (panel B) extracted from an event related task design. Data were collected with an anterior to posterior phase encoding direction (7 Tesla, TR 1.4s, Multiband 2, iPAT 3, 6/8ths Partial Fourier, 0.8 mm isotropic resolution, TE 27.4ms, Flip Angle 78°, Bandwidth 1190Hz) and covered the occipital pole and ventral temporal cortex. Six runs of data were collected. Stimuli consisted of faces *versus* fully phase scrambled variants of the same faces. Stimuli were presented for 2 seconds, with a 2 second interstimulus interval. A. After smoothing with a 0.8mm FWHM Gaussian kernel, the maps (middle) show larger areas of activation, however, at a cost of spatial precision compared to NORDIC (right). Note that areas associated with narrow or small areas of activation are not present in smoothed data compared to NORDIC (Inset, left area). In addition, the boundary between positive and negative t-statistics is less sharp compared to NORDIC, reflecting a reduction of spatial precision due to smoothing (inset, right area). B. To summarize the effects, t-statistics for various methods were extracted using an ROI produced from 6 runs of the Standard reconstruction (blue) contrasting faces vs. scrambled. NORDIC (orange) produces t-statistic values that exceed the Standard dataset (blue). NORDIC t-statistics is more comparable to, but slightly better than that observed when the original data is spatially smoothed with a 0.8mm FWHM gaussian filter (red); however, NORDIC achieves the improvement in t-statistics without spatially blurring the image.

Spatial filtering achieves improvements in t-statistics by averaging signals over regions that are larger than those determined by the voxel dimensions set by the acquisition parameters, therefore greatly compromising the spatial resolution of the data. NORDIC instead can be thought of as having the same positive effect on the ability to detect stimulus/task induced signal changes in fMRI as spatial filtering *<u>without</u>* the deleterious consequences of image blurring that come with spatial filtering of images.

#### iii) fMRI with AUDITORY STIMULUS

As a demonstration of yet another different sensory stimulus and different cortical region, we present data obtained with auditory stimuli. The results again are comparable to those presented in the main text as well as the additional data included in this supplementary material as shown in Supplementary Figs. 8 through 11. Namely, multiple runs obtained over significantly longer data acquisition times are needed with Standard reconstruction to achieve the extent of activation comparable to a single NORDIC run.

**Supplementary Figure 12:**
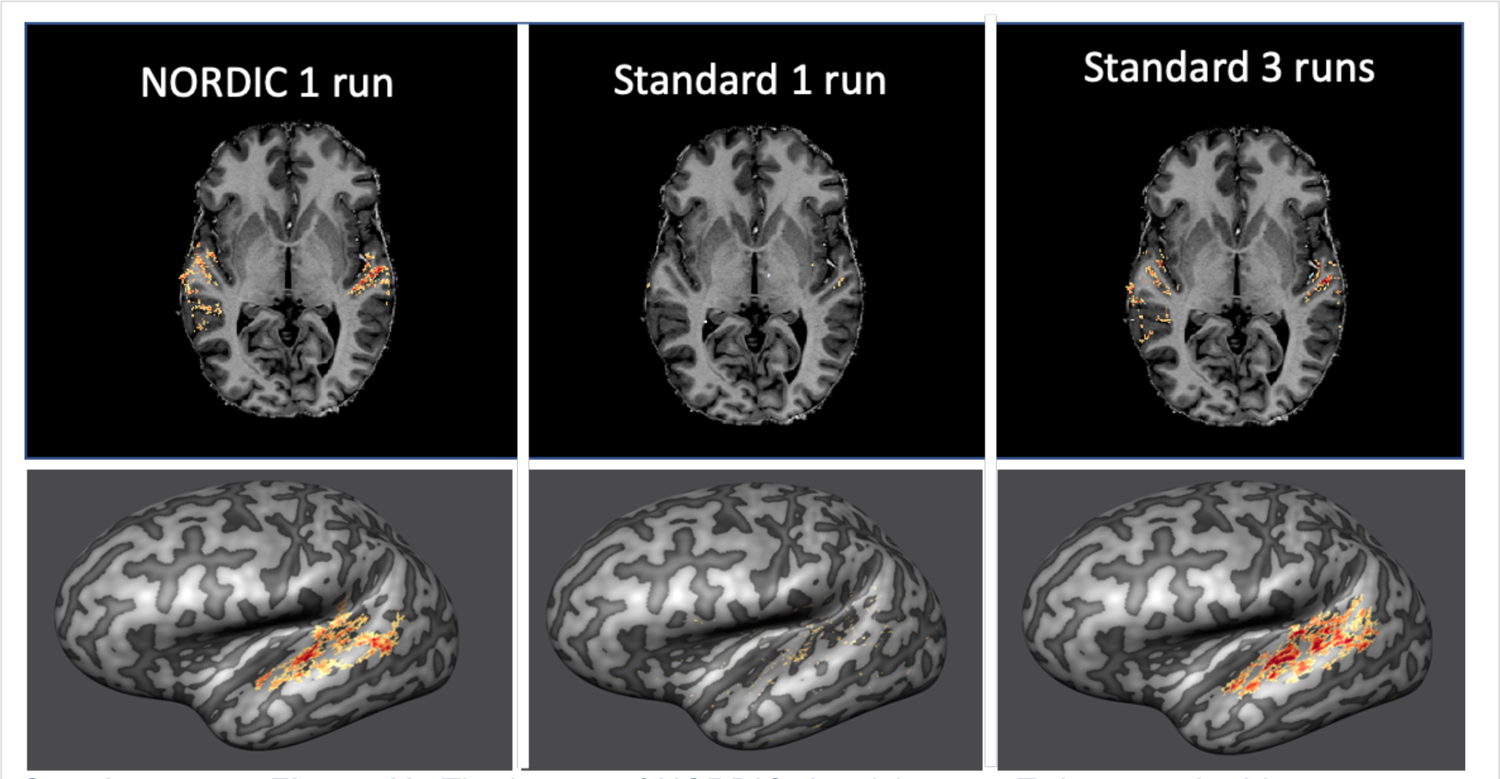
The impact of NORDIC denoising on 7T data acquired in response to sound presentation (0.8 mm iso voxels; TR 1600 ms; MB 2; iPAT 3; 42 slices). Sounds (simple tones) were presented following a slow event related design (each sound lasting 1 second were presented with an average inter trial interval of 10 seconds) while participants passively listened to them. The figure shows t-maps for the main effect of sound presentation (i.e. sound vs. silence; thresholded at t > 2.7) in volume space (top row – one representative transversal slice) and in inflated cortical space (bottom row – left hemisphere). Note, 3 Standard runs (approximately 18 minutes of data) are required to achieve the extent of activation comparable to a single NORDIC run (approximately 6 minutes of data). These results highlight advantages conferred by NORDIC denoising beyond application to visual experiments.

#### *iv)* Resting state fMRI

The bulk of this work focused on task/stimulus based fMRI to demonstrate in great detail the impact of NORDIC on fMRI because in many ways task fMRI provides the ability to calculate numerous parameters, such as percent signal change, t-statistics of detection of task/stimulus induced signal changes, functional point spread function etc. that can be quantitatively evaluated for the consequences of NORDIC processing. However, resting-state fMRI (rsfMRI) is also a frequently employed approach in neuroscience research, and is a major component of the HCP^1, 2^. The NORDIC technique should work for rsfMRI. Here, we provide a preliminary demonstration of the beneficial impact of NORDIC on resting state data.

The resting state acquisition consisted of four 10 minute runs with 3T HCP acquisition parameters (Supplementary Figure 13). No stimulus presentation occurred and participants were instructed to stay still, minimize movements, and fixate on a visible crosshair.

Minimal processing steps, performed with AFNI, were applied to the Standard and NORDIC data. These included slice timing correction and motion correction to the first volume of the first run of the Standard data for both Standard and NORDIC data. Next, we regressed out the 6 estimates of motion parameters and polynomials up to 5^th^ order. A spherical seed, with radius of 3mm was placed in the medial prefrontal cortex, corresponding to a location within the Default Mode Network. The extracted seed time course for each run was used to generate a map of Pearson’s r values, corresponding to the correlation of each voxel in the brain with the seed timeseries (i.e. seed-based correlation).

These preliminary resting state NORDIC results are indeed promising. However a more in dept look at resting state acquisition and processing is required to fully appreciate the impact of NORDIC denoising on these types of acquisitions and analyses.

**Supplementary Figure 13:**
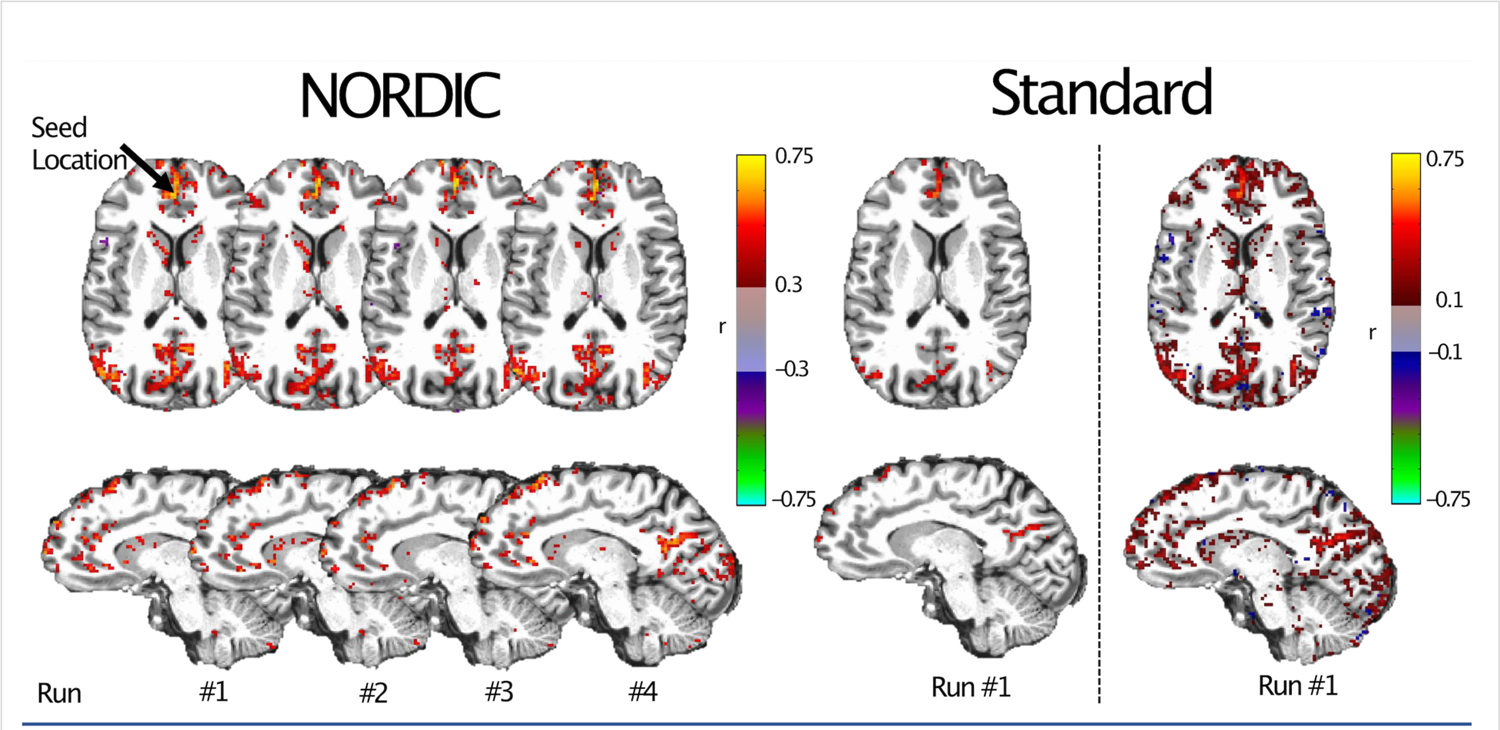
Resting State fMRI. NORDIC processing benefits for resting state, seed-based connectivity. Correlation maps, with Pearson’s r values are shown for 4 sequential runs of resting state data. The seed time course was extracted from the medial prefrontal cortex, corresponding to a node within the Default Mode Network. Correlation maps obtained individually from each 10 min run are highly consistent between the 4 runs of the NORDIC processed data (Left), and show cortical and subcortical correlations with r > 0.3 that are not visible in the Standard data at the same threshold (Middle). Only by reducing the standard data threshold to r>0.1 do similar patterns emerge, albeit with more punctate and scattered apparent correlations including in the white matter (Right most panel).

#### ***v)*** 0.5 mm isotropic data

**Supplementary Figure 14:**
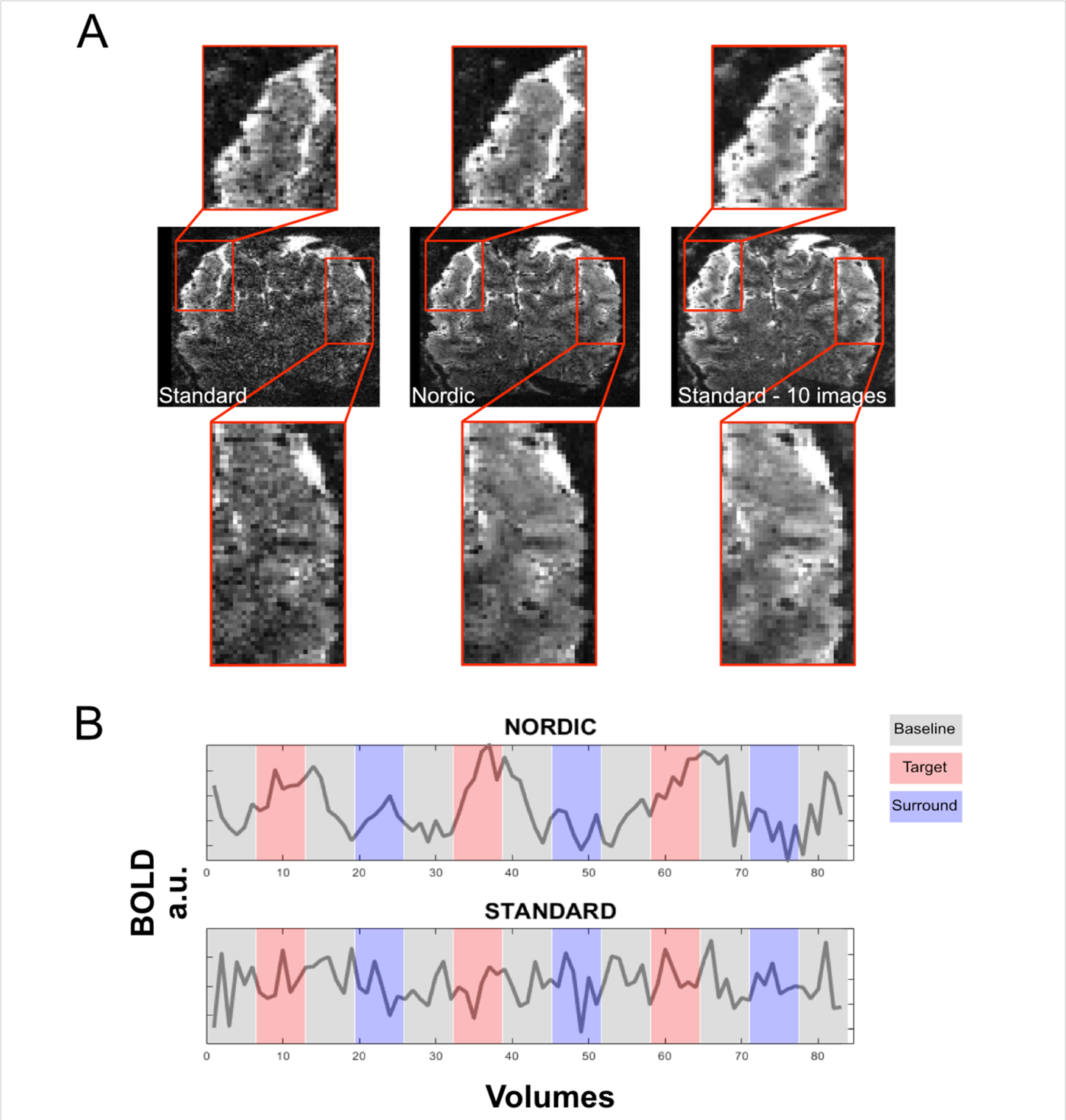
Additional features of the 0.5 mm isotropic resolution data shown in Figure 6 of the main manuscript. Panel A shows a single slice from a single time point in the consecutively acquired volumes forming the fMRI time series, processed with Standrd reconstruction (left-most), NORDIC (middle) and sum of 10 images of Standard reconstruction (right most). Zoomed-in inserts demonstrate that fine features of the image observed in Standard reconstructed images are preserved in NORDIC, consistent with lack of blurring. Panel B shows the timecourse of a single medial voxel for NORDIC (top) and Standard (bottom) reconstructions froom a single run.The data were acquired using a 3D gradient echo (GE) EPI approach with a spatial resolution of 0.5 mm isotropic voxels (3D GE EPI; 40 slices; iPAT 3; TR 83 ms; Volume acqusition time 3652 ms). The total data acqusition time for a single run fMRI time series was ∼ 5 minutes (i.e. 2 visual conditions - center and surround gratings – each presented 3 times for 24 seconds, and each followed by a 24 seconds fixation period.

**Supplemental Figure 15.**
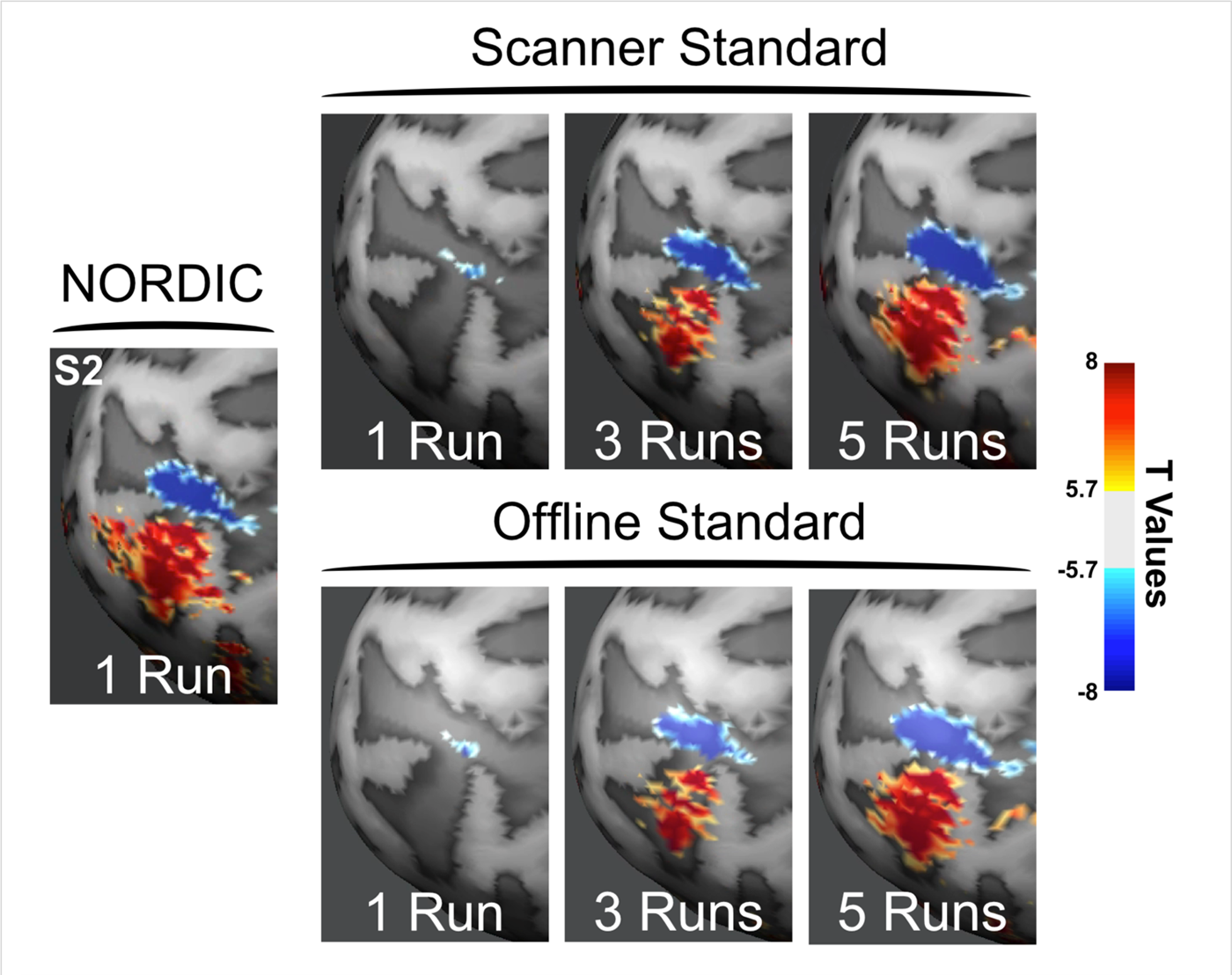
Comparing NORDIC to “Offline” and “Scanner” standard. Top row of 3 images labelled “Scanner Standard” is a duplicate of Figure 2 for subject S2 in the main body of the paper; these functional maps, displayed for a single run and for the concatenation of 3 or 5 runs, were derived from fMRI time series of accelerated GE EPI images generated using scanner image reconstruction from the acquired k-space data. For the lower row of corresponding images labeled “Offline Standard”, the same k-space data were exported offline and were processed with our offline pipeline including EPI and GRAPPA reconstructions but *without* the NORDIC denoising step. It can be seen that the two different Standard reconstructions, Scanner *versus* Offline Standard, produce virtually identical results, and for both Standards, it takes ∼5 concatenated runs to achieve equivalence to NORDIC denoised single run. All functional maps and derivative results in the main body of the paper compare NORDIC denoised results, generated by our “offline” pipeline, against those derived from fMRI time series generated using the scanner (Siemens, 7T) provided image reconstruction. It is, therefore, important to demonstrate that the scanner reconstruction produce comparable results to our offline *in the absence of* NORDIC denoising. Otherwise, the differences seen with NORDIC can be ascribed not only to the NORDIC approach but also differences in other aspects of image reconstruction. Conversely, had we only used our offline pipeline both for standard and denoised images, one can argue that our EPI and GRAPPA reconstruction may be inferior and the gains we attribute to NORDIC actually would be less had we simply used the scanner reconstruction. The results presented in this figure demonstrate that such arguments are not valid, and that without NORDIC, the scanner and our offline reconstruction produces virtually identical results.

**Supplemental Figure 16:**
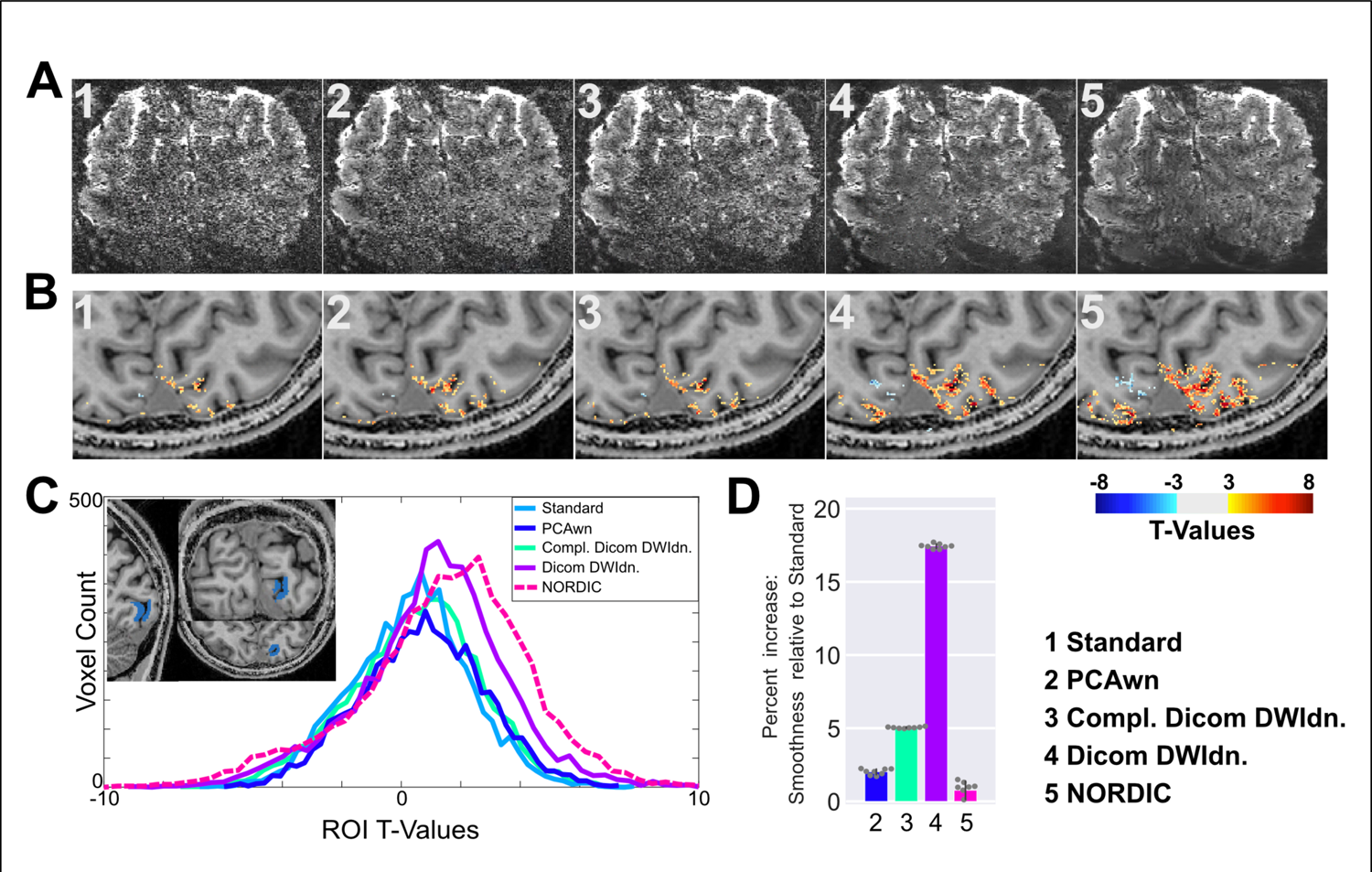
Impact of different denoising strategies on 0.5 mm isotropic resolution functional data. The top row (panel A) shows a representative slice from GE EPI images acquired for different reconstructions: (1) Standard (scanner reconstruction); (2) global PCA denoising with white noise criteria (PCAwn) as described in refernce^3^ to identify random noise; (3) DWI Denoise method performed on complex dicoms (Compl.DicomDWIdn); (4) DWI Denoise performed on magnitude only dicoms (Dicom DWIdn); and (5) NORDIC. The second row (panel B) shows the t-thresholded functional maps (t>|3|) computed for the contrast target > surround performed on the 8 concatenated runs for the same 5 reconstructions. Panel C shows, the t-value distributions for the same target>surround contrast for these 5 reconstructions within an ROI (shown in the top-left inlay) hand drawn on the co-registered T1wigthed images at the approximate location of the retinotopic representation of the target in V1. Panel D shows the smoothness metrics for all denoising methods in percent increase relative to the scanner Standard. For all 8 runs, smoothness was computed independently per run and estimated with a spatial autocorrelation metric using a Gaussian+monoexponential decay model as in Figure 5C in the main manuscript (see Methods for more details). Error bars represent standard errors of the mean across the 8 runs. As discussed in the results section of the main manuscript, NORDIC shows the best performance, i.e. the largest right shift for t value distribution (dashed red line) with virtually no smoothing. The t-distributions for Standard, PCAwn, and Compl. DICOM DWIdn are highly comparable; even though a relatively small improvement, accompanied by a small increase in smoothness, can be appreciated for the t-values computed for PCAwn, and Compl. DICOM DWIdn relative to Standard. The second best performance as judged only by the t-value distribution is from Dicom DWIdn; however, this is achieved with significant smoothing, which alone could be responsible for part of the improved performance. These observations are also reflected in the functional maps and the individual EPI images. *Taken together*, the metrics presented here are useful in evaluating the performance of different denoising algorithm. However, caution should be exercised in interpreting anyone metric alone, as discussed at the end of the Results section in the main manuscript.

